# Convergence of distinct subpopulations of mechanosensory neurons onto a neural circuit that elicits grooming

**DOI:** 10.1101/2020.06.08.141341

**Authors:** Stefanie Hampel, Katharina Eichler, Daichi Yamada, Hyunsoo Kim, Mihoko Horigome, Romain Franconville, Davi D. Bock, Azusa Kamikouchi, Andrew M. Seeds

## Abstract

Diverse subpopulations of mechanosensory neurons detect different mechanical forces and influence behavior. How these subpopulations connect with central circuits to influence behavior remains an important area of study. We previously discovered a neural circuit that elicits grooming of the *Drosophila melanogaster* antennae that is activated by an antennal mechanosensory chordotonal organ, the Johnston’s organ (JO) (Hampel et al., 2015). Here, we describe anatomically and physiologically distinct JO mechanosensory neuron subpopulations and define how they interface with the circuit that elicits antennal grooming. We show that the subpopulations project to distinct zones in the brain and differ in their responses to mechanical stimulation of the antennae. Each subpopulation elicits grooming through direct synaptic connections with a single interneuron in the circuit, the dendrites of which span the different mechanosensory afferent projection zones. Thus, distinct JO subpopulations converge onto the same neural circuit to elicit a common behavioral response.

## Introduction

Animals can detect complex mechanical forces in their environments through an array of different mechanosensory neuron types that respond to specific stimuli. These neurons influence appropriate behavioral responses through their connections with circuits in the central nervous system (CNS). One prominent feature of mechanosensory systems as they project into the CNS from the periphery is the topographical arrangement of their axonal projections in distinct zones. For example, mammalian auditory neurons can project to specific zones based on their excitation by particular vibrational frequencies (Appler and Goodrich, 2011; Kandler et al., 2009). Thus, subpopulations that are similarly tuned project to the same zone in the CNS, and the collective projection zones of subpopulations that are tuned to different frequencies form a tonotopic map. Different types of topographical organization are described across mechanosensory neuron types that are based on tonotopy or anatomical location (somatotopy). Yet it remains unclear how sensory topography interfaces with neural circuits to influence behavior (Kaas, 1997; Thivierge and Marcus, 2007).

The fruit fly (*Drosophila melanogaster*) Johnston’s organ (JO) is a chordotonal organ in the antennae that can be studied to address this outstanding question. The JO detects diverse types of mechanical forces that move the antennae, including sound, wind, gravity, wing beats, and tactile stimuli (Hampel et al., 2015; Ishikawa et al., 2017; Kamikouchi et al., 2009; Mamiya and Dickinson, 2015; Matsuo et al., 2014; Patella and Wilson, 2018; Yorozu et al., 2009). The ability of the JO to respond to these different stimuli is conferred by about 480 mechanosensory neurons called JONs (Kamikouchi et al., 2006). Subpopulations of JONs are selectively excited by different vibrational frequencies or by static displacements of the antennae and send their projections into discrete zones in the CNS (Kamikouchi et al., 2006). In accordance with their diverse physiological tuning properties, the JONs are implicated in controlling diverse behaviors including courtship, locomotion, gravitaxis, escape, flight, and grooming (Hampel et al., 2015; Kamikouchi et al., 2009; Lehnert et al., 2013; Mamiya and Dickinson, 2015; Mamiya et al., 2011; Tootoonian et al., 2012; Vaughan et al., 2014; Yorozu et al., 2009). The recent development of genetic tools in *Drosophila* has enabled the identification and experimental manipulation of downstream neural circuitry that controls some of these behaviors (Hampel and Seeds, 2017), including sound responses (e.g. courtship song) (Kamikouchi et al., 2009; Lai et al., 2012; Vaughan et al., 2014), escape (Matsuo et al., 2016; Pézier et al., 2014), and grooming (Hampel et al., 2015). Other studies are using these tools to define how JON-mediated mechanosensory responses are processed through these neurons (Azevedo and Wilson, 2017; Chang et al., 2016; Lehnert et al., 2013; Patella and Wilson, 2018; Tootoonian et al., 2012). However, the precise synaptic connectivity of the different JON subpopulations with downstream neurons remains largely unknown. A recently produced dataset that consists of a complete serial section electron microscopy (EM) volume of the adult fruit fly brain now enables high-resolution mapping of all neurons and their synaptic connections (Zheng et al., 2018). This dataset was used to define how two distinct JON subpopulations are synaptically connected with neurons involved in escape behavior and courtship song responses (Kim et al., 2020). The studies described above provide the foundation for studies that seek to define the circuit mechanisms by which subpopulations of JONs interface with downstream circuits to control behavior.

We previously discovered that activation of different JON subpopulations elicits antennal grooming behavior (Hampel et al., 2015), which involves the grasping and brushing of the antennae by the front legs (Böröczky et al., 2013; Robinson, 1996). At least two different subpopulations of JONs were found to elicit grooming, however we did not determine the extent to which these subpopulations were morphologically and physiologically distinct from each other. We also found that the JONs elicit grooming by activating a neural circuit that elicits or “commands” grooming of the antennae, and comprises three different morphologically distinct interneuron types. Two types are located where the JON projections terminate in a region of the ventral brain called the subesophageal zone (SEZ), and were named antennal grooming brain interneurons 1 and 2 (aBN1 and aBN2). The third type includes a cluster of descending neurons (aDNs) that have their dendrites in the SEZ and axonal projections in the ventral nerve cord (VNC). The aDNs are the proposed outputs of the circuit because they project to the region of the VNC where the antennal grooming pattern generation circuitry is presumed to be located (Berkowitz and Laurent, 1996; Burrows, 1996). The different JON subpopulations that elicit grooming are in close proximity with these different interneuron types, and at least one subpopulation is functionally connected with the circuit. This suggested that different JON subpopulations converge onto the circuit to control grooming behavior. However, the synaptic connectivity of the JONs with the different interneurons in the circuit was not defined.

Here, we describe how the JON subpopulations that elicit grooming are distinct from each other and define how they converge onto the antennal grooming command circuit. Because the subpopulations are each known to consist of morphologically heterogeneous JON types that have not been completely described (Kamikouchi et al., 2006), we first define their morphological diversity by reconstructing major portions of each subpopulation from an EM volume of the fly brain (Zheng et al., 2018). We next produce transgenic driver lines that selectively target expression in each subpopulation. These lines enable us to visualize the distribution of the different subpopulations in the antennae and determine that they respond differently to mechanical stimuli. Optogenetic activation experiments confirm our previous finding that each JON subpopulation can elicit grooming of the antennae (Hampel et al., 2015). However, we report here that one of the subpopulations also elicits an avoidance response, whereas the other elicits wing flapping movements. Finally, by employing EM reconstructions and functional imaging experiments, we determine that JONs within each subpopulation are synaptically and functionally connected with the antennal command circuit through the same interneuron (aBN1). These results provide a comprehensive description of the topography of the JO, and reveal how different JON subpopulations whose projections occupy different points in topographical space can converge on the same neural circuit to elicit a similar behavioral response.

## Results and discussion

### Electron microscopy-based reconstruction of different JON subpopulations

We first sought to define the morphological diversity of the neurons within each JON subpopulation as a prerequisite for understanding how the subpopulations interface with the antennal command circuit. JONs project from the antennae through the antennal nerve into a region of the SEZ called the antennal mechanosensory and motor center (AMMC) (**Figure 1A**). The projections of different subpopulations form distinct zones in the AMMC (zones A-E, **Figure 1B**). JONs that respond to antennal vibrations project laterally into zones A and B (called JO-A and -B neurons), whereas JONs that are tuned to antennal vibrations and/or static deflections project medially into zones C-E (JO-C, - D, and -E neurons) (Kamikouchi et al., 2009; Mamiya and Dickinson, 2015; Matsuo et al., 2014; Patella and Wilson, 2018; Yorozu et al., 2009). We previously discovered a subpopulation that we refer to here as projecting to zone F (JO-F neurons, referred to as aJO neurons in our previous work). JO-F neurons enter the brain through the AMMC like other JONs, but then project ventrally (**Figure 1A,B**, blue JONs) (Hampel et al., 2015). Because our previous work implicated JO-C, -E, and -F neurons in grooming behavior, we reconstructed these subpopulations within a serial-section EM volume of the entire fruit fly brain to define their morphological diversity (Zheng et al., 2018).

**Figure 1.**
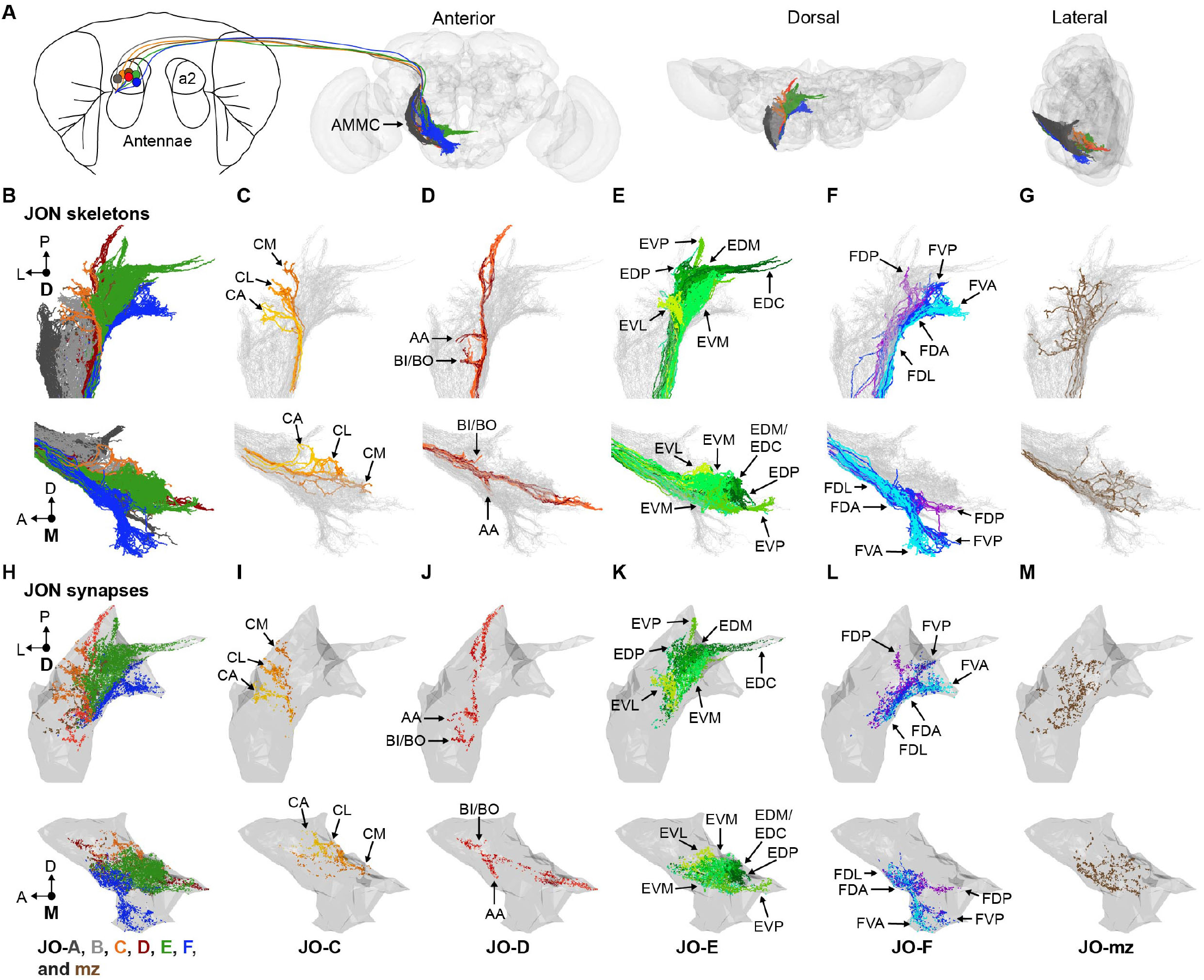
EM-based reconstruction of JONs. (**A**) JON projections from the second segment of the antennae (a2) into the AMMC brain region (outline of brain neuropile is shown in gray). Anterior, dorsal, and lateral views of the reconstructed JONs are shown. (**B-G**) Reconstructed JONs are shown from the dorsal (top) and medial (bottom) views. (**H-M**) Dorsal (top) and medial (bottom) views of the JON pre- and postsynaptic sites are shown (colored dots) with a gray mesh that outlines the entire reconstructed JON population. All reconstructed JONs are shown in **B**, but only fully reconstructed JONs are shown in **H** (JO-A and -B synapses not shown). JO-A and -B neurons were previously reconstructed (Kim et al., 2020). Colors in **A**, **B**, and **H** correspond to the zones to which the different JONs project, including zones A (dark gray), B (light gray), C (orange), D (red), E (green), F (blue), and mz (brown). Panels **C-M** show JONs that project specifically to zones C (**C,I**), D (**D,J**), E (**E,K**), F (**F,L**), or multiple zones (mz) (**G,M**). Color shades in **CM** indicate different neuron types that project to that zone. Zone subareas are indicated with labeled arrows.

We first located the JO-C, -E, and -F neurons in the EM volume. A confocal z-stack of a driver line (R27H08-GAL4) expressing GFP in these JON subpopulations was registered into the EM volume (**Figure 1 – figure supplement 1A,B**) (Bogovic et al., 2018; Hampel et al., 2015; Zheng et al., 2018). We then examined the JON axon bundle where the antennal nerve enters the brain and found that the GFP had highlighted the medial region of the bundle where the JO-C and -E neurons were previously described to reside (Kamikouchi et al., 2006). Lateral to this region were the previously reconstructed JO-A and -B neurons (Kim et al., 2020). We reconstructed 147 JONs within the GFP-highlighted region (**Figure 1 – figure supplement 1C**). 104 were completely reconstructed, including all of their pre- and postsynaptic sites. The remaining 43 JONs were reconstructed using an autosegmentation algorithm that identified the main branches, but not finer branches or synapses (Li et al., 2019). These JONs were useful for examining gross morphology, but not for determining connectivity.

JON topography can be defined based on the segregated organization of the different zones in the AMMC, and the stereotyped projections of the JONs to distinct subareas in each zone. We therefore compared previous light microscopy-based descriptions of these two features with our EM reconstructed JONs to categorize them into specific zones (**Figure 1B-G**, **Video 1**). This not only confirmed that we had reconstructed JO-C, -E, and -F neurons (**Figure 1C,E,F**), but revealed that we had also reconstructed JO-D neurons, as well as JONs that project to multiple zones (JO-mz neurons, **Figure 1D,G**).

Three lines of evidence suggested that the reconstructed JONs represent the major diversity of the JO-C, -D, -E, and -F neurons. First, when the reconstructed JONs were viewed *in toto*, we could observe each previously described subarea (**Figure 1B-F**). This suggested that we had not missed JONs that are major contributors to these subareas. Second, in comparison with previous estimates of the numbers of JO-C, -D, and -E neurons (Kamikouchi et al., 2006), we have reconstructed major portions of each. Based on estimates obtained from stochastic labeling experiments, we have reconstructed 29% of the JO-C neurons (9 out of an estimated 31 neurons), 69% of the JO-D neurons (9 out of 13), and 87% of the JO-E neurons (62 out of 71). However, counts of JO-CE neurons targeted by particular transgenic driver lines suggest that there are almost twice the number of JO-CE neurons. Therefore, the calculated percentages of reconstructed JONs may be overestimates. Third, and as we show below, the JONs could be categorized based on their morphological similarities to each other. The fact that we reconstructed multiple morphologically similar JONs suggests that we captured the diversity of each subpopulation. However, it is possible that reconstruction of more JONs would uncover additional diversity, especially among the JO-mz neurons, of which we only reconstructed 7 out of an estimated 58 neurons (12%). These reconstructions provide the most comprehensive description of the JO-C, -D, -E, -F, and -mz neurons to date. In combination with the previously reconstructed JO-A and -B neurons (Kim et al., 2020), we now provide a near complete description of the diversity of JONs that make up each subpopulation in the JO (**Figure 1B**). This will provide a valuable resource for studies seeking to understand the neural circuit basis of the JO chordotonal organ’s functions.

### The topographical organization of different JON subpopulations

The EM reconstructions enabled us to next systematically identify the morphologically distinct JON types within each subpopulation. Below we provide a nomenclature for each JON type and then define how they contribute to the topographical organization of each zone (see **Video 1** for 3D overview, see **Supplementary file 1** for a full list of reconstructed JONs).

The lateralmost nine of the reconstructed JONs were JO-C neurons whose projections form three different subareas (**Figure 1C**). Two were previously named zone C medial (CM) and lateral (CL) (Kamikouchi et al., 2006). The third was a previously undescribed subarea located anterior of CM and CL (named C anterior (CA)). By examining the projections of individual reconstructed JO-C neurons, we found that each subarea is mainly formed by one of three JO-C neuron types. Two of these types project exclusively to a single subarea to form either CL or CA. However, the third type whose projections form CM has a smaller branch that projects to CL. Based on these observations, we named the three JO-C neuron types according to the subarea that receives their largest branch (named JO-CM, -CL, and -CA neurons, **Figure 1C**, **Figure 1 – figure supplement 2A-C**).

Nine of the reconstructed JONs project to zone D (**Figure 1D**). The proximal region of zone D contains protrusions that extend towards either zones A or B, and were previously named AA and BI/BO, respectively (Kamikouchi et al., 2006). Two different JO-D neuron types were previously described and named JO-D posterior (JO-DP) and JO-D anterior (JO-DA) neurons (**Figure 1D**, **Figure 1 – figure supplement 3A,B**). The JO-DA neurons have branches that extend to both AA and BI/BO, and then a projection that extends towards, but does not reach the posteriormost region of zone D. The JO-DP neurons tend to have fewer second-order branches than JO-DA neurons, and extend a projection to the posteriormost region of the zone.

62 of the reconstructed JONs project to zone E. This zone has five previously described subareas that we identified from the reconstructed JONs (**Figure 1E**) (Kamikouchi et al., 2006). The subareas are formed when JO-E neurons enter the brain and then split into two adjacent bundles called E dorsomedial (EDM) and E ventromedial (EVM). EDM curves medially and approaches the midline to form the E dorsal in the commissure (EDC) subarea. JONs that form EDC were named JO-EDC neurons, whereas most of the other JONs terminate earlier in the EDM bundle and were named JO-EDM neurons **(Figure 1 – figure supplement 4A,B**). Another JON type in EDM forms a posterior protrusion called the dorsoposterior (EDP) subarea (named JO-EDP neurons). JONs within the other major bundle, EVM, were divided into three types and named JO-EVM, -EVP, and -EVL neurons (**Figure 1 – figure supplement 4B**). JO-EVM neurons remain in the EVM bundle. JO-EVP neurons form a protrusion from EVM that projects to the posterior brain called the E ventroposterior (EVP) subarea. JO-EVL neurons form a newly described subarea called E ventrolateral (EVL) that projects laterally from EVM, towards zone C. Some of the reconstructed JONs were not morphologically similar to the other JO-E neurons. The branches of these JONs tiled the ventralmost region of zone E, and were therefore named zone E ventral (JO-EV) neurons.

60 of the reconstructed JONs project to zone F and form five subareas (**Figure 1F**). The first three are formed by the proximal neurites of JO-F neurons that branch to different parts in the AMMC. We named these subareas zone F dorsoanterior (FDA), dorsoposterior (FDP), and dorsolateral (FDL). FDA is formed by JO-F neurons that extend a branch that runs adjacent to the JO-EVM neurons. Lateral and slightly ventral to FDA is the relatively small anterior protruding FDL subarea. Some JO-F neurons form the FDP subarea by extending a posterior branch that projects adjacent to the JO-EVP neurons (**Figure 1E,F**). The distal neurites of JO-F neurons project to the ventral SEZ in two bundles that form the ventroanterior (FVA) and ventroposterior (FVP) subareas (**Figure 1F**). Five JO-F neuron types form the different zone F subareas (**Figure 1F**, **Figure 1 – figure supplement 5A,B**). The first type that we named JO-FVA neurons contain few or no second-order branches and project through the AMMC to the ventral SEZ, where most terminate their projections in the FVA subarea. The second type that we named JO-FDA neurons project to FDA in the AMMC, and then ventrally to FVA and/or FVP. The third type that we named JO-FDP neurons project to FDA and FDP in the AMMC and then ventrally to the FVP subarea. The last two types that were named JO-FDL and -FVL neurons both project to the FDL subarea. These types differ in that the JO-FDL neurons terminate dorsally in the FDL subarea, whereas the JO-FVL neurons also have a projection in the ventral SEZ.

In contrast to the JONs that project to specific zones, we reconstructed seven JO-mz neurons that have branches projecting to multiple zones (**Figure 1G**). These JONs have been previously identified (Kamikouchi et al., 2006), but little is known about their morphological diversity, functional roles, or tuning properties. Six out of seven reconstructed JO-mz neurons have branches that follow the posterior projections of the JO-FDP neurons, while extending projections to other zones (**Figure 1G**, **Figure 1 – figure supplement 6A,B**). We could not classify the JO-mz neurons into subtypes because they showed no clear morphological similarity.

The above described topographical organization was based on the projections of the JONs. We next addressed the extent to which this organization was reflected at the synaptic level. Previous immunohistochemical experiments indicated that JON presynaptic sites are broadly distributed in the posterior regions of each zone (Kamikouchi et al., 2006). We found that the synapses of the 104 completely reconstructed JONs were also distributed throughout the posterior regions of their respective zones (**Figure 1H-M**). Because these synapses included both pre- and postsynaptic sites (**Supplementary file 1**), we compared their relative distributions in each zone. Examination of the total distribution of synapses did not reveal any clear difference, as the subareas of each zone had both pre- and postsynaptic sites (**Figure 1 – figure supplement 7A-F**). Taken together, our results show that different JON types project in a segregated manner, and form discrete zones and subareas that contain both pre- and postsynaptic sites. Thus, the subareas interface with downstream circuits, but are likely also subject to regulation.

### NBLAST clustering of the reconstructed JONs

Our reconstructions enabled us to categorize the JONs based on their morphology and projections into specific zones and subareas. Because this was done by manual annotation, we used the NBLAST clustering algorithm as an independent categorization method (Costa et al., 2016). NBLAST uses spatial location and neuronal morphology to calculate similarity between neurons. In agreement with our manual annotations, NBLAST clustered many of the same JONs that we had assigned as specific types, such as JO-EVL or -EVP neurons (**Figure 2A,B**, groups 3 and 14, **Supplementary file 1** shows the JON types and their NBLAST group number). NBLAST also revealed that manually assigned JON types could be further subdivided. For example, the algorithm divided JO-EVM neurons into two clusters that occupied distinct regions in the EVM subarea (**Figure 2A,B**, groups 5 and 6). JO-mz neurons were not all clustered together, consistent with our annotation of these neurons as projecting to different zones. In some cases, NBLAST clustered JONs that we had assigned as distinct from each other. For example, some JO-FVA and -FDA neurons were clustered into the same group (**Figure 2A,B**, groups 9 and 10). These differences likely arose based on relatively small branches from the main projections of these JONs that were differentially emphasized by our annotations versus the NBLAST algorithm. Despite these differences, our annotations and NBLAST both show how a topographical map is formed by different JON subpopulations that project to distinct zones and subareas.

**Figure 2.**
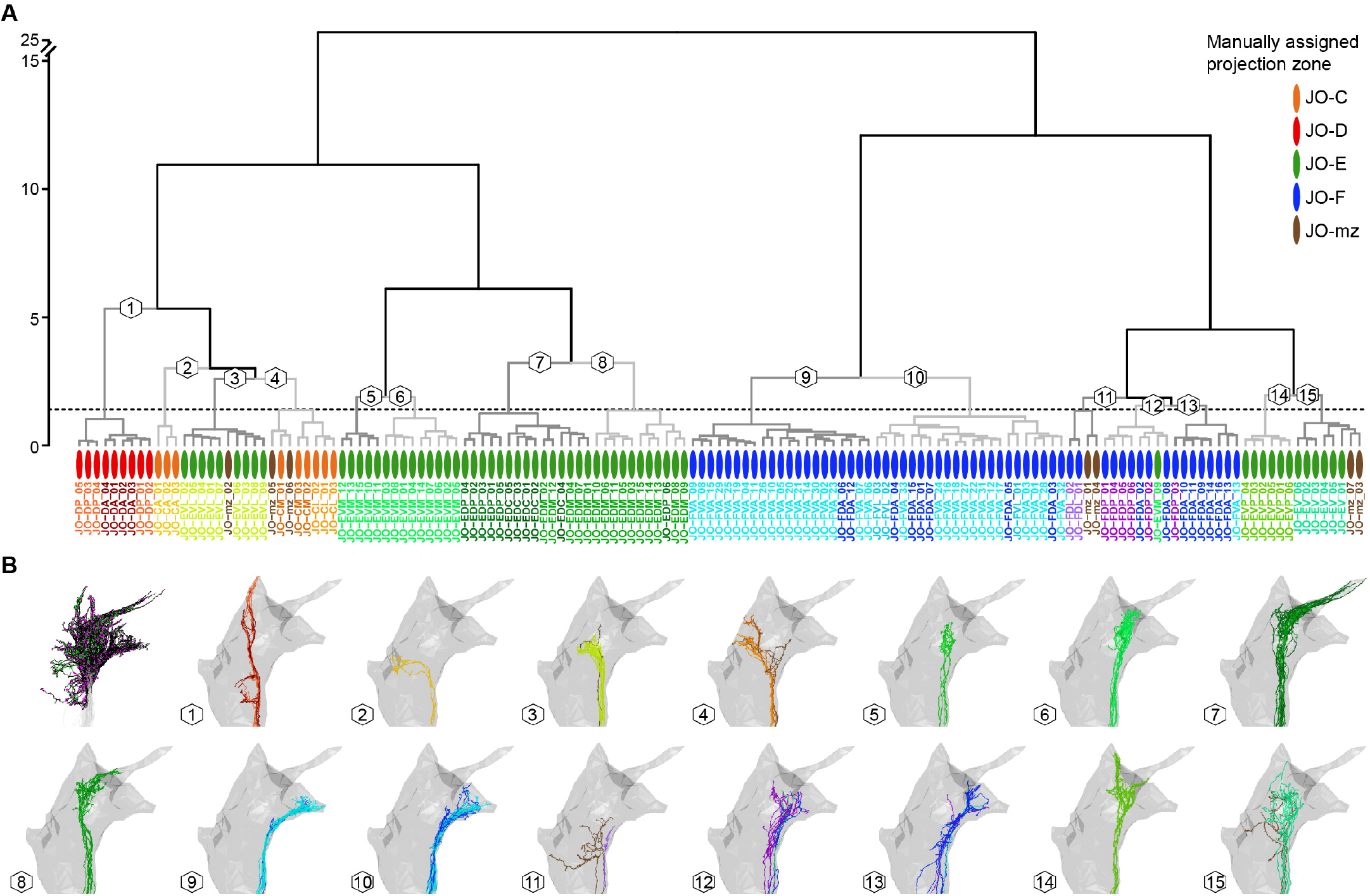
Correspondence between manual annotation and NBLAST clustering in the categorization of JONs. (**A**) Dendrogram of hierarchically clustered scores from an NBLAST query of 147 reconstructed JONs. Oval colors indicate the manual asignment of each JON as projecting to a specific zone. Colors of specific neuron names indicate the manually annotated neuron types (e.g. JO-FVA in cyan, JO-FDA in dark blue). Branch numbers indicate JON groups that are shown in **B** resulting from a cut height of 1.4 (dotted line). (**B**) NBLAST clustered JON groups (15 groups at h = 1.4). Neuron colors indicate manually annotated neuron types and correspond to the neuron type names in **A**. The first panel shows how the JONs were pruned for NBLAST analysis to include only synapse bearing parts of the neurons (JON parts used for NBLAST analysis shown in black, see Materials and methods for details) with pre- and postsynaptic sites in magenta and green respectively.

### Driver lines that express in JO-CE or -F neurons

We next produced transgenic driver lines that would enable us to compare the anatomical, physiological, and behavioral properties of the JO-CE and -F neurons. New lines were necessary because there were no previously reported drivers that expressed exclusively in JO-C and -E neurons. Further, although we previously described a “clean” line that expresses in JO-F neurons (aJO-spGAL4-1) (Hampel et al., 2015), here we obtained additional lines to expand our toolkit for genetically accessing these JONs. We used a Split GAL4 screening approach to produce four different lines that expressed in JONs whose activation could elicit antennal grooming (see Materials and methods for details, **Figure 3 – figure supplement 1A-D**). Two of the identified lines express in JO-C and -E neurons and were named JO-CE-1 (spGAL4 combination: 100C03-AD ⋂ R27H08-DBD) and JO-CE-2 (R39H04-AD ⋂ R27H08-DBD) (**Figure 3A,B**). The other two lines express mainly in JO-F neurons and were named JO-F-1 (R25F11-AD ⋂ R27H08-DBD) and JO-F-2 (122A08-AD ⋂ R27H08-DBD) (**Figure 3C,D**). Our analysis of all four driver lines revealed no evidence of JO-A, -B, - D, or -mz neurons in their expression patterns.

**Figure 3.**
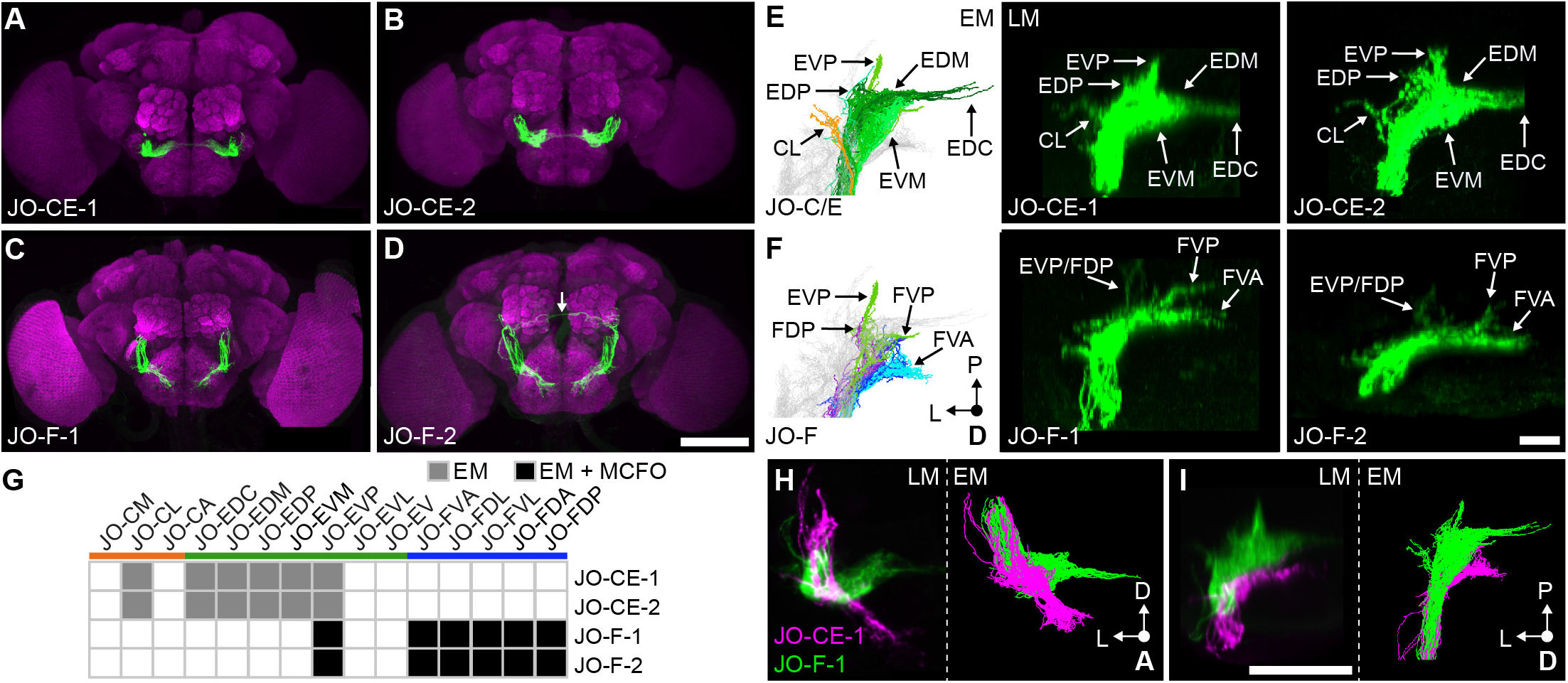
Driver lines that express in JO-CE or JO-F neurons. (**A-D**) Shown are maximum intensity projections of the brains (anterior view) of JO-CE-1 (**A**), JO-CE-2 (**B**), JO-F-1 (**C**), and JO-F-2 (**D**) expressing green fluorescent protein (mCD::GFP). The brains are co-stained with anti-GFP (green) and anti-Bruchpilot (magenta). Scale bar, 100 μm. The arrow shown in **D** indicates a neuron that is not a JON. (**E,F**) EM reconstructed JON types (left panels) that are predicted to be in the expression patterns of each line shown in the middle and right panels (dorsal view). Expression patterns of JO-CE-1 (middle) and −2 (right) are shown in **E**, and JO-F-1 (middle) and −2 (right) are shown in **F**. Subareas are indicated with arrows. Note that in **F** the subareas FDL and FDA are not labeled because they are not visible in the dorsal view. Scale bar, 20 μm. (**G**) Table of JON types that are proposed to be in each expression pattern. The shade of each box indicates whether the predictions are supported by EM reconstructions alone (gray), or by EM and MCFO data (black). (**H,I**) Computationally aligned expression patterns of JO-CE-1 (magenta) and JO-F-1 (green) from anterior (**H**) and dorsal (**I**) views (left panels) in comparison with the EM reconstructed JONs (right panels). Scale bar, 50 μm.

The EM reconstructed JONs were next used to assess which JON types were targeted by JO-CE-1 and −2. The subareas formed by the reconstructed JONs (**Figure 3E**, left) were compared with those observed in the confocal images of each driver line expressing mCD8::GFP (**Figure 3E**, middle and right). Both lines express in JONs projecting to the subareas CL, EDP, EVP, EDM, EDC, and EVM. Because each subarea is formed by specific JON types, we could deduce that both lines express in JO-CL, -EDC, -EDM, -EDP, -EVM, and -EVP neurons (**Figure 3G**). In contrast, we could not identify the CM, CA, or EVL subareas in the expression patterns of either line, suggesting that JO-CE-1 and −2 do not express in JO-CM, -CA, and -EVL neurons. It was unclear if the lines express in JO-EV neurons, as these JONs do not form a specific subarea that would aid in their identification. We sought to verify the different JON types in each expression pattern using the multicolor flipout (MCFO) method to stochastically label individual JONs (Nern et al., 2015). However, we were unable to label individual JONs using this method with JO-CE-1 or −2. Importantly, neither line expresses in JO-F neurons as there are no ventral-projecting JONs in their patterns (**Figure 3A,B**). We concluded that JO-CE-1 and −2 express specifically in JO-C and -E neurons (**Figure 3G**).

JO-F-1 and −2 express in JONs projecting to each zone F subarea, including FDA, FDP, FDL, FVA, and FVP (**Figure 3F**). Based on the EM-defined JON types that make up each subarea, we proposed that both lines express in JO-FVA, -FDL, -FVL, -FDA, and -FDP neurons (**Figure 3G**). It was unclear if the lines also expressed in JO-E neurons because of the possibility that these JONs were obscured by the JO-FDA and -FDP neurons. Therefore, we used the MCFO method to label individual JONs within each pattern. Using this method, we could observe the JO-FDA, -FDP, -FDL, - FVL, and -FVA neurons, with the majority of them being JO-FDA neurons (**Figure 3 – figure supplement 2A-E**). A portion of the labeled JONs projected to zone E and had a posterior projection, leading us to propose they are JO-EVP neurons (**Figure 3 – figure supplement 2F**). However, the lines only weakly labeled the EVP subarea as compared with the JO-F subareas (**Figure 3E,F**), suggesting that a relatively small number of JO-EVP neurons are labeled. We concluded that JO-F-1 and −2 express mostly in JO-F neurons, but also in JO-EVP neurons. Of note, JO-F-1 and −2 appear to express in the same JON types as our previously reported JO-F driver line named aJO-spGAL4-1 (Hampel et al., 2015).

To visualize the extent to which the JO-CE and -F driver lines express in distinct JON subpopulations, we computationally aligned confocal stacks of their expression patterns in the same brain (**Figure 3H,I**, left panels). This shows how the different driver lines express in JON subpopulations with projections into distinct zones. Further, the morphology of the aligned projections was very similar to the EM reconstructed JO-CE and -F neurons (**Figure 3H,I**, right panels). This provides further support that the different lines selectively target the JO-CE or -F neurons.

We compared the distributions of the JONs that are labeled by the different driver lines in the JO chordotonal organ. The JON cell bodies are organized into a bottomless bowl-shaped array in the second antennal segment that can be visualized by labeling the JON nuclei using an antibody against the ELAV protein (**Figure 4A,B**). Expression of GFP under control of JO-CE-1 and −2 labeled JON cell bodies in a ring around the JO bowl (**Figure 4C,D**). In contrast to the previously published JO-CE drivers that showed expression around the entire ring of the JO bowl (Kamikouchi et al., 2006), JO-CE-1 and −2 showed only sparse expression around the anterior dorsal (A-D) portion of the bowl (**Figure 4C’,D’**). JO-F-1 and −2 showed expression in two clusters in the dorsal and ventral regions of the JO bowl (**Figure 4E,F**), in agreement with our previous results (Hampel et al., 2015). The dorsal expression was in the anterior and posterior regions of the bowl (A-D and P-D), while the ventral expression was largely restricted to the posterior (P-V) region (**Figure 4E’,F’**). In contrast to what we previously reported, JO-F-1 and −2 expression was not restricted to the dorsal and ventral clusters, but was also in more intermediate JONs in the posterior part of the bowl. This prompted us to reexamine JO-F driver lines from our previous work for evidence that they also expressed in these intermediate JONs (Hampel et al., 2015). Indeed, these lines show relatively faint GFP signal in intermediate JONs in the posterior bowl (not shown). This suggests that the distribution of JONs targeted by the different JO-F driver lines is more continuous, rather than restricted to clusters. A comparison of the distributions of JONs that are targeted by the JO-CE and -F driver lines revealed that they occupy common (P-D and P-V) and distinct regions (A-D and A-V) of the JO bowl (**Figure 4B’-F’**). However, the functional significance of these JON distributions remains unclear.

**Figure 4.**
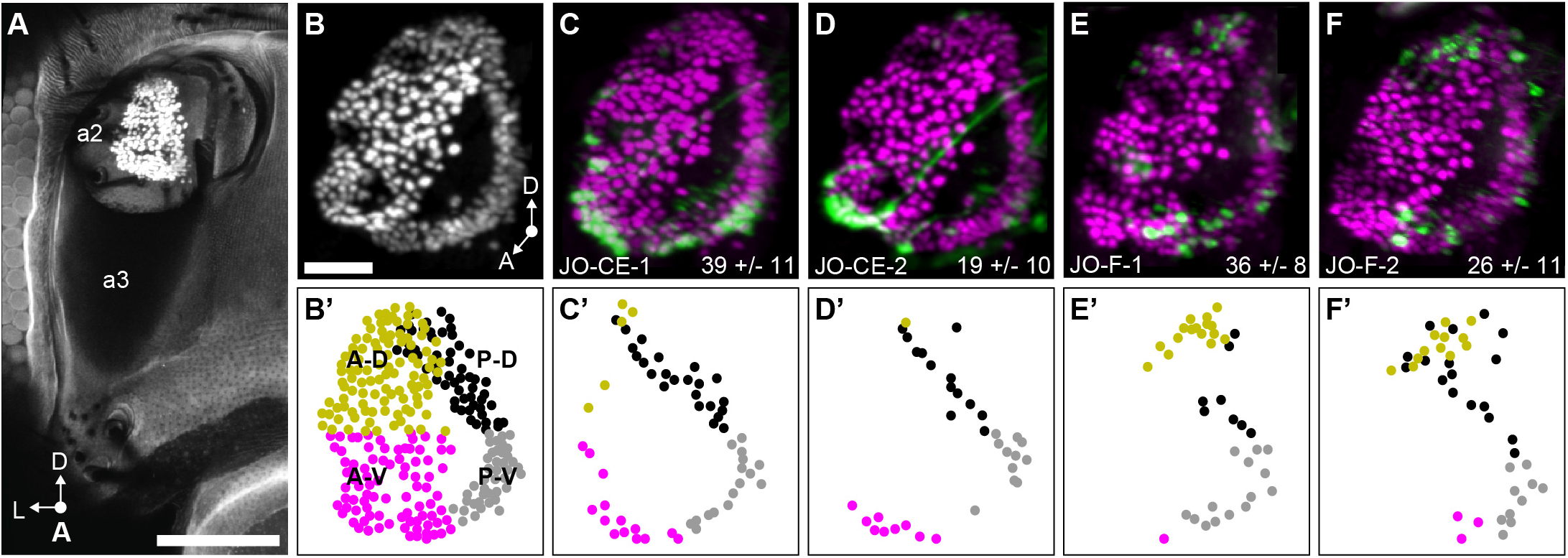
Distribution in the JO chordotonal organ of JONs that are targeted by JO-CE and F driver lines. (**A**) Anterior view of the antennal region of the head with the JON nuclei labeled with an anti-ELAV antibody in the second antennal segment (labeled a2, third segment is labeled a3). A maximum intensity projection is shown. The head is visualized as autofluorescence from the cuticle. Scale bar, 100 μm. (**B**) JON nuclei laterally rotated about the ventral/dorsal axis (~40°). Scale bar, 25 μm. (**C-F**) Driver lines expressing GFP in the JO. Shown is a co-stain with anti-GFP (green) and anti-ELAV (magenta) antibodies. The average number of JONs labeled in each line ± the standard deviation is shown in the bottom right corner. (**B’**) Anterior view of manually labeled JON cell bodies in different regions from the confocal stack shown in **B** (not laterally rotated like in **A-F**, ~60% JON cell bodies labeled). The JO regions are color coded, including anterior-dorsal (A-D, mustard), posterior-dorsal (P-D, black), anterior-ventral (A-V, magenta), and posterior-ventral (P-V, gray). (**C’-F’**) Manually labeled JON cell bodies (dots) that expressed GFP in a confocal z-stack of each driver line. This highlights GFP-labeled JONs in the posterior JO that are difficult to view in the maximum projections shown in **C-F**. The colors indicate the JO region where the cell body is located. Shown are JO-CE-1 (**C,C’**), JO-CE-2 (**D,D’**), JO-F-1 (**E,E’**), and JO-F-2 (**F,F’**).

### JO-CE and -F neurons respond differently to stimulations of the antennae

We next compared the responses of the JO-CE and -F neurons to mechanical stimulations of the antennae using a previously established preparation (Matsuo et al., 2014). Flies expressing the calcium indicator GCaMP6f (Chen et al., 2013) in the JONs targeted by the different driver lines were immobilized and their mouthparts removed to obtain optical access to the JON axon terminals in the brain. Stimuli were delivered using an electrode to move the arista and third antennal segment (**Figure 5A**). The rotation of the third segment about the second segment in a particular direction or sinusoidal frequency excites the JONs. Thus, we imaged calcium responses in the JON axons in the brain while different stimuli were applied.

**Figure 5.**
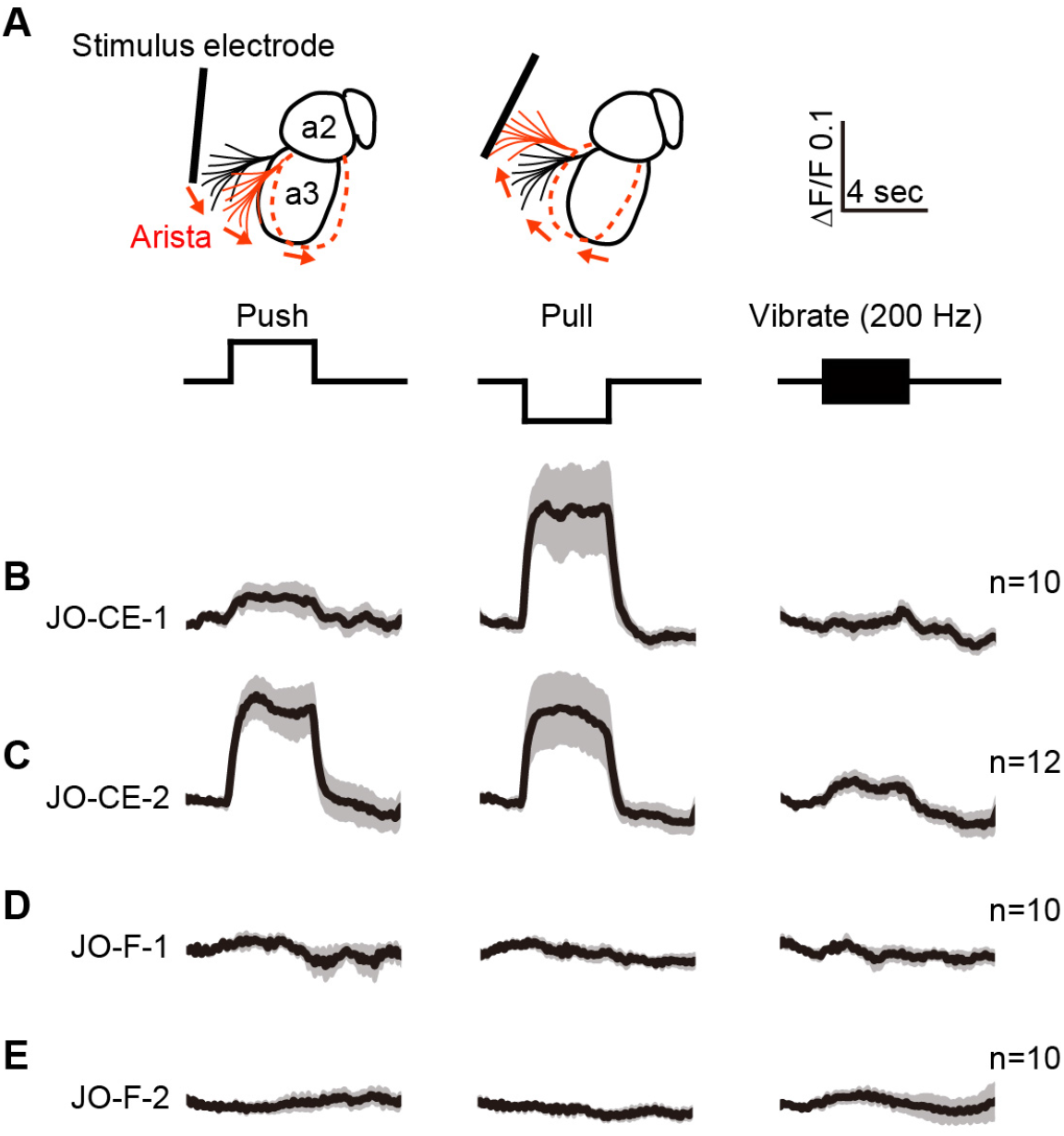
Testing the responses of JO-CE and JO-F neurons to stimulations of the antennae. (**A**) Schematic lateral view of a fly antenna. An electrostatically charged electrode pushes or pulls the antenna via the arista towards or away from the head, respectively, or induces a 200 Hz sinusoid. (**B-C**) Calcium response of JONs to stimulations of the antennae. Flies were attached to an imaging plate, dorsal side up. The proboscis was removed to access the ventral brain for imaging GCaMP6f fluorescence changes (ΔF/F) in the JON afferents. Stimulations of the antennae were delivered for 4 seconds as indicated above the traces. Each row shows the mean trace (black lines) from a different driver line expressing GCaMP6f, including JO-CE-1 (**B**), JO-CE-2 (**C**), JO-F-1 (**D**), JO-F-2 (**E**). The gray envelopes indicate the standard error of the mean. n ≥ 10 for each line per stimulus.

JO-CE neurons were previously found to respond to static deflections that either push or pull the antennae towards or away from the head (Kamikouchi et al., 2009; Patella and Wilson, 2018; Yorozu et al., 2009). In accordance with this finding, the JONs labeled by JO-CE-1 and −2 showed increased GCaMP6f fluorescence in response to both push and pull of the arista (**Figure 5B,C**, **Figure 5 – figure supplement 1A-D**). Also in agreement with previous results using this immobilized fly preparation (Kamikouchi et al., 2009; Matsuo et al., 2014; Yorozu et al., 2009), we found no evidence that the JO-CE neurons responded to vibrations (**Figure 5B,C**, **Figure 5 – figure supplement 1A-D**, 200 Hz test shown). However, other work has shown that the JO-CE neurons can respond to wing beat-generated vibrations while flies are flying (Mamiya and Dickinson, 2015). This may indicate that JO-CE neurons are tuned to vibrations during flight, and to static pushes and pulls of the antennae in other behavioral states.

It is unknown what stimulus excites JO-F neurons. Under the experimental conditions used here, the JONs that are labeled by JO-F-1 or −2 did not respond to push or pull movements of the antennae (**Figure 5D,E**, **Figure 5 – figure supplement 2A-D**). Furthermore, we could not identify a vibration frequency that could evoke a response in these JONs, including low (40 Hz, n=2 flies), middle (200 Hz, N=10 flies), and high frequency vibrations (400 and 800 Hz, n=2 flies) (**Figure 5D,E**, **Figure 5 – figure supplement 2A-D**, 200 Hz test shown). We confirmed that the JONs were competent to respond to stimuli by applying KCl at the end of each experiment and observing an increased GCaMP6f signal (not shown). This may suggest that a novel stimulus excites these JONs. However, it is also possible that a response to one of the tested stimuli can be observed under different experimental conditions, such as in a walking or flying fly. This could be tested using an experimental preparation that enables flies to walk or fly while being imaged (Mamiya and Dickinson, 2015). However, our results here indicate that JO-CE and -F neurons do not show similar responses to mechanical stimuli in immobilized flies. Thus, the JO-CE and -F neurons are both anatomically and physiologically distinct from each other.

### Activation of JO-CE or JO-F neurons elicits common and distinct behavioral responses

We next compared the extent to which the JO-CE and -F neurons influence common and distinct behaviors. Our previous work implicated these subpopulations in eliciting the common behavior of antennal grooming (Hampel et al., 2015). While no other behaviors have been ascribed to the JO-F neurons, the JO-CE neurons are implicated in such behaviors as wind-induced suppression of locomotion, gravitaxis, and the modulation of flight (Kamikouchi et al., 2009; Mamiya and Dickinson, 2015; Yorozu et al., 2009). In the present study, we compared the breadth of overt behavioral changes that are caused by activating either JO-CE or -F neurons. The red light-gated neural activator CsChrimson (Klapoetke et al., 2014) was expressed using the different JO-CE and JO-F driver lines. Flies were placed in chambers so that they could move freely and then exposed to red light for optogenetic activation of the JONs (Hampel et al., 2015, 2017).

We first reproduced our previous results by showing that activation of either JO-CE or -F neurons elicits grooming (**Figure 6A**, **Videos 2 and 3**). However, the JO-F-1 and −2 driver lines express in JO-F and -EVP neurons, which raised the possibility that the JO-EVP neurons were responsible for the grooming rather than the JO-F neurons (**Figure 3G**). To address this possibility, we identified another driver line named JO-F-3 (R60E02-LexA) that expresses exclusively in JO-F neurons whose activation elicited grooming (**Figure 6 – figure supplement 1A-G**). Thus, the data presented here further implicate the JO-CE and -F neurons in antennal grooming. However, it remains unclear whether the JO-C and -E subpopulations are both responsible for the grooming. Addressing this will require obtaining transgenic driver lines that express exclusively in one subpopulation or another.

**Figure 6.**
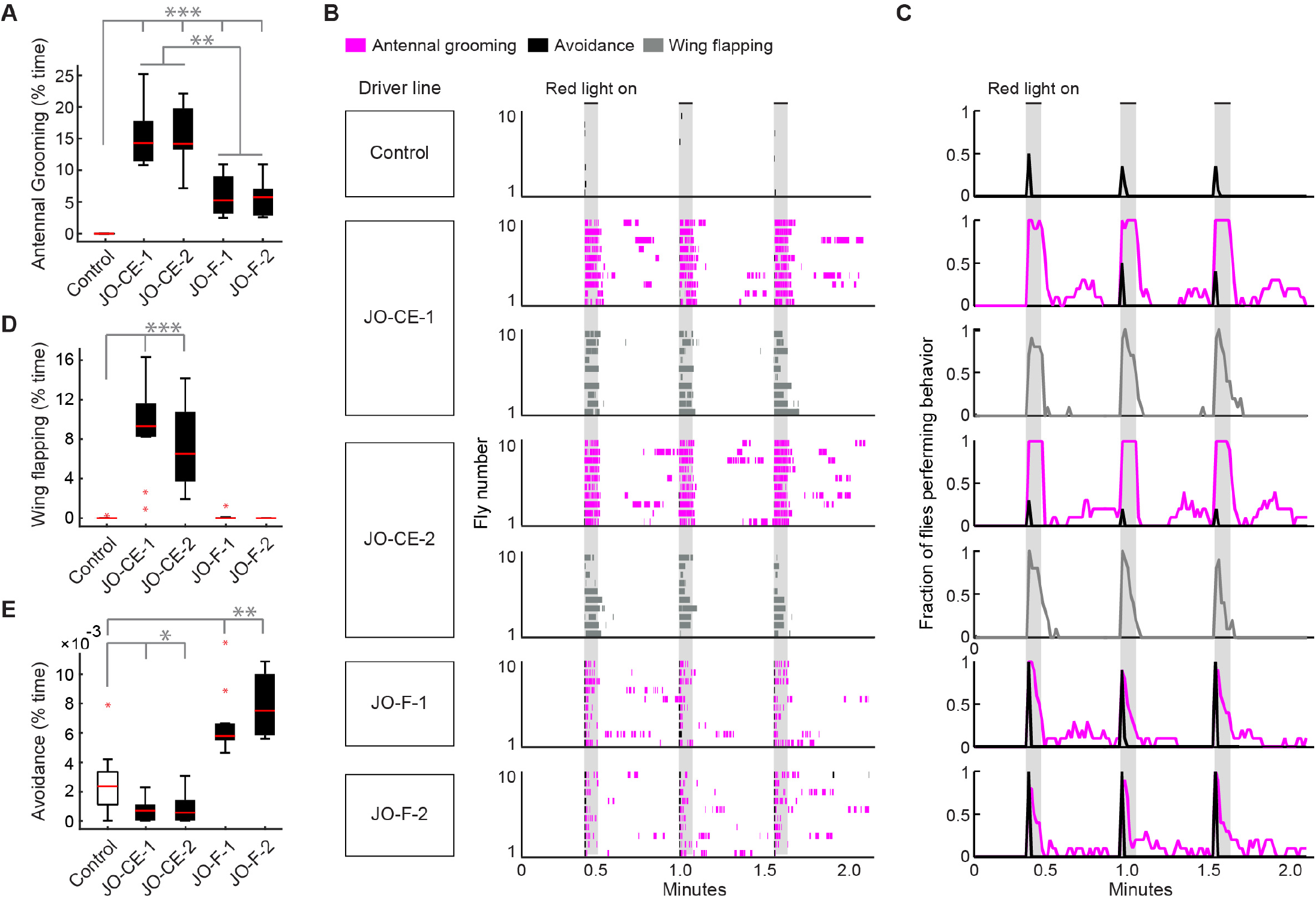
Optogenetic activation of either JO-CE or JO-F neurons elicits distinct behavioral responses. (**A, D, E**) Percent time flies spent performing antennal grooming (**A**), wing flapping (**D**), or avoidance (**E**) with optogenetic activation of JONs targeted by JO-CE-1, JO-CE-2, JO-F-1, and JO-F-2. Control flies do not express CsChrimson in JONs. Bottom and top of the boxes indicate the first and third quartiles respectively; median is the red line; whiskers show the upper and lower 1.5 IQR; red dots are data outliers. n ≥ 10 for each box; asterisks indicate *p < 0.05, **p < 0.001, ***p < 0.0001, Kruskal–Wallis and post hoc Mann–Whitney U pairwise tests with Bonferroni correction. (**B**) Ethograms of manually scored video show the behaviors elicited with red light induced optogenetic activation. Ethograms of individual flies are stacked on top of each other. The behaviors performed are indicated in different colors, including antennal grooming (magenta), wing flapping (gray), and avoidance (black). Light gray bars indicate the period where a red light stimulus was delivered (5 sec). (**C**) Histograms show the fraction of flies that performed each behavior in one-second time bins. Note that only JO-CE-1 and −2 elicited wing flapping, which was not mutually exclusive with grooming. Therefore, an extra row of wing flapping ethograms and histograms is shown for those lines.

This experiment further revealed that the JO-CE and -F neurons elicit grooming that lasts for distinct durations after the onset of the optogenetic stimulus (**Figure 6A-C**, **Figure 6 – figure supplement 1E,F**, magenta traces). Five second optogenetic activation of the JO-CE neurons elicited grooming that lasted throughout the duration of the stimulus. In contrast, the JO-F neurons elicited shorter duration grooming that terminated prior to the stimulus cessation. We considered the trivial possibility that these distinct durations of grooming were caused by differences in the number of activated JONs that were targeted in each line. However, the average number of labeled JONs did not differ markedly between the JO-CE and -F driver lines (**Figure 4C-F**). This suggests that the distinct grooming durations were due to the physiological properties and/or functional circuit connectivity of each subpopulation.

In the process of annotating the grooming performed by flies with optogenetic activation of the JO-CE neurons, we observed the flies simultaneously performing wing flapping movements (**Figure 6B-D**, JO-CE-1 and −2, gray trace, **Video 2**). The wings would extend to approximately 45-90-degree angles from the body axis while flapping. This observation is intriguing given that the JO-CE neurons were previously shown to modulate wing movements during flight (Mamiya and Dickinson, 2015). Our finding that wing flapping results from optogenetic activation of the JO-CE neurons provides new evidence of that these JONs can modulate the wing movements.

Activation of JO-F neurons elicited a backwards locomotor response that appeared as if flies were avoiding an object that bumped into their antennae (**Video 3, Figure 6B,C**, **Figure 6 – figure supplement 1H**). The avoidance and grooming were mutually exclusive, as the avoidance occurred briefly at the onset of the stimulus and was immediately followed by grooming (**Figure 6B,C**, JO-F-1 and −2, black and magenta traces). Control flies also show avoidance in response to the red light stimulus (**Figure 6B,C**, control, black trace, **Video 4**). However, less than half of these flies responded (42%), whereas nearly all of the JO-F neuron activation trials showed avoidance (97% for JO-F-1, 100% for JO-F-2). JO-F neuron activation also elicited longer lasting avoidance responses than controls, with the experimental flies spending between 5 and 8-fold more time in avoidance behavior than control flies (**Figure 6E**). These results implicate locomotor avoidance as a behavior that is stimulated by the JO. In stick insects, a backwards locomotor response is elicited by mechanical stimulations of the antennae (Graham and Epstein, 1985), however, the mechanoreceptor(s) that mediate this response are unknown. Our results may suggest JO-F neurons as a link between mechanical stimulations of the antennae and locomotor avoidance.

### EM reconstruction of the antennal command circuit brain interneurons aBN1 and aBN2

The JO-CE and -F neurons elicit common and distinct behaviors, with antennal grooming as the common behavior and wing flapping and avoidance as the distinct behaviors. We hypothesized a circuit organization that could explain this, whereby the JON subpopulations have converging inputs onto the antennal command circuit and diverging inputs onto putative circuits that control either wing flapping or avoidance (**Figure 7A**). In the case of avoidance, two interneuron types (MAN and MDN) were previously identified that elicit backwards locomotion (Bidaye et al., 2014). MDN was also found to be necessary for a vision-based avoidance response, revealing that these neurons can respond to sensory inputs (Sen et al., 2017; Wu et al., 2016). Thus, the JO-F neurons could elicit an avoidance response through functional connections with MAN/MDN-like neurons. The JO-CE neurons are proposed to impinge on the wing motor system through descending circuitry (Mamiya and Dickinson, 2015), however the neurons in this pathway remain to be identified. Here, we sought to test whether the JO-CE and -F neurons have converging inputs onto the antennal command circuit.

**Figure 7.**
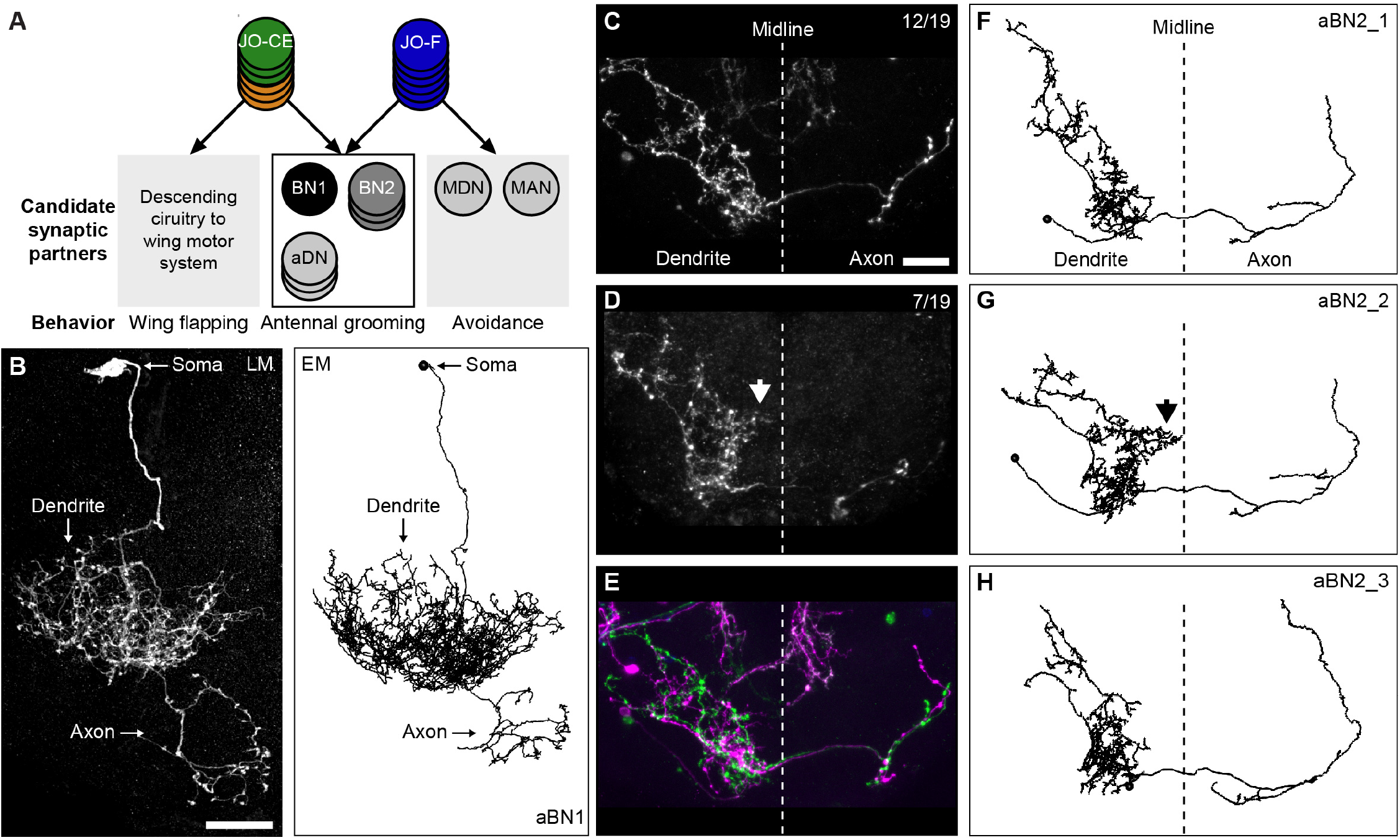
EM reconstruction of antennal command circuit interneurons aBN1 and aBN2. (**A**) Three putative circuits are proposed to receive converging or diverging inputs from the JO-CE and -F neurons as indicated with arrows. In the gray boxes are candidate synaptic partners that could elicit each behavior. Labeled circles indicate previously identified interneuron candidates (Bidaye et al., 2014; Hampel et al., 2015). Black box indicates the circuit that is the focus of this study. (**B**) Light microscopy-imaged (LM, left) and EM-reconstructed (EM, right) aBN1 neuron. The dendrite, axon, and soma are indicated with labeled arrows. Scale bar, 20 μm. (**C,D**) MCFO labeling of individual aBN2 neurons in the aBN2-spGAL4-1expression pattern. Shown are anterior views of maximum intensity projections of two different aBN2 neurons that were stained using a tag-specific antibody (see Materials and methods for information about the tagged proteins and antibodies used). The aBN2 neuron shown in **D** has a distinct midline projecting branch that is indicated with an arrowhead. Top right corner of each panel indicates the number of individually labeled aBN2 neurons without (**C**) or with (**D**) the midline projecting branch, versus the total MCFO-labeled aBN2 neurons. Approximate brain midline is indicated with a dotted line. Scale bar, 20 μm. (**E**) MCFO labeling of two aBN2 neurons in the same brain hemisphere that both lacked the midline-projecting branch. The green neuron is also shown in **C**. (**F-H**) The three EM reconstructed aBN2 neurons (names are indicated in the top right corner). Neurons are shown as skeletons without synapses. Approximate brain midline is indicated with a dotted line. The midline projecting branch in aBN2_2 (**G**) is indicated with an arrowhead.

We hypothesized that the JO-CE and -F neurons interface with the command circuit through connections with one or both of its brain interneuron types (i.e. aBN1 and aBN2, **Figure 7A**). This was based on four lines of evidence from our previous work (Hampel et al., 2015). First, optogenetic activation of either of these neuron types is sufficient to elicit antennal grooming. Second, JO-C, -E, and -F neuron-elicited grooming is disrupted by silencing either aBN1 or aBN2. Third, activation of the JO-C, -E, and -F neurons induces calcium responses in aBN1 and aBN2, demonstrating that they are functionally connected. Fourth, the JONs and the aBNs are all in close proximity in the AMMC and ventral brain. We therefore focused our EM reconstructions on aBN1 and aBN2 in this work. A future study will examine the synaptic connectivity of the descending neurons (aDNs), which are functionally connected with the JONs, aBN1, and aBN2.

We began to search for aBN1 and aBN2 in the EM volume by tracing postsynaptic partners of a reconstructed JO-EDM neuron (JO_EDM_07, **Figure 1 – figure supplement 4B**). Rather surprisingly, the first postsynaptic neuron that we identified was aBN1. This was confirmed by comparing the neuron’s morphology with the confocal-imaged neuron that was described in our previous work (Hampel et al., 2015). Both the light-microscopy-imaged and the EM-reconstructed aBN1 had a highly branched dendritic field in the AMMC (**Figure 7B**, dendrite). Both had a branched axon that extended into the ventromedial SEZ (**Figure 7B**, axon). Furthermore, while the dendritic and axonal branches were located in the ventral brain, the soma was uniquely located in the dorsal brain (**Figure 7B**, soma). Our previous experiments suggested that there was only a single aBN1 neuron that was necessary for JO-stimulated antennal grooming (Hampel et al., 2015). Consistent with this, our searches in the EM volume only identified one neuron (not shown). aBN1 was completely reconstructed, including all of its synapses.

Our previous work indicated that there were three aBN2 neurons (Hampel et al., 2015). This was based on the observation that the aBN2 driver line used in that study, aBN2-spGAL4-1, labeled a cluster of three neurons whose silencing disrupted the grooming response to JON activation. We presumed that these neurons were morphologically similar, however did not show this. In order to identify and reconstruct these aBN2 neurons from the EM volume, we first defined their morphology using the MCFO method to individually label each neuron that is targeted by aBN2-spGAL4-1. All neurons that were labeled using this method were morphologically similar, with an ipsilateral dendritic field and an axon that projects to the contralateral brain hemisphere (**Figure 7C,D**, two examples shown). The branched dendrite extends from the AMMC to the ventromedial SEZ. 7 out of 19 (37%) individually labeled aBN2 neurons were observed to have a dendritic branch that approaches the midline of the brain (**Figure 7D**, arrowhead). We propose that one of the three aBN2 neurons has this branch given that we observed labeling of two aBN2 neurons in the same brain hemisphere that both lacked the midline-projecting branch (**Figure 7E**).

We next identified and reconstructed the aBN2 neurons in the EM volume. Our limited searches of postsynaptic partners of different JONs did not identify these neurons, leading us to search for aBN2 as a putative postsynaptic partner of aBN1 (Hampel et al., 2015). We indeed identified a neuron that was morphologically similar to the MCFO-labeled aBN2s (named aBN2_1, **Figure 7F**). Given that our light-microscopy data indicated that the aBN2 axons project through the brain as a bundle (example shown in **Figure 7E**), we searched around the axon of aBN2_1 and identified two additional morphologically similar neurons (named aBN2_2 and aBN2_3, **Figure 7G,H**). Consistent with our previous data, we only identified three aBN2-like neurons in our searches through the EM volume (not shown). Further, one of these aBN2 neurons (aBN2_2) had a midline-projecting dendritic branch, in agreement with what we had predicted based on MCFO data (**Figure 7G**). All three neurons were fully reconstructed including their synapses.

### Different JON subpopulations have converging inputs onto the antennal command circuit

The complete reconstruction of aBN1 and aBN2 neurons enabled us to next define their connectivity with the JONs. The reconstructed JON, aBN1, and aBN2 neurons were all in close proximity in the ventral brain (**Figure 8A,B**), in agreement with what we previously showed using computationally aligned confocal stacks of these neurons (Hampel et al., 2015). We therefore analyzed the all-to-all connectivity among these different neurons (major connections shown in **Figure 8C** and **Figure 8 – figure supplement 1**, all-to-all connectivity shown in **Supplementary file 2**). Our analysis included the previously reconstructed JO-A and -B neurons (Kim et al., 2020), which enabled us to define the connectivity of aBN1 and aBN2 neurons with a representative portion of all subpopulations that make up the JO.

**Figure 8.**
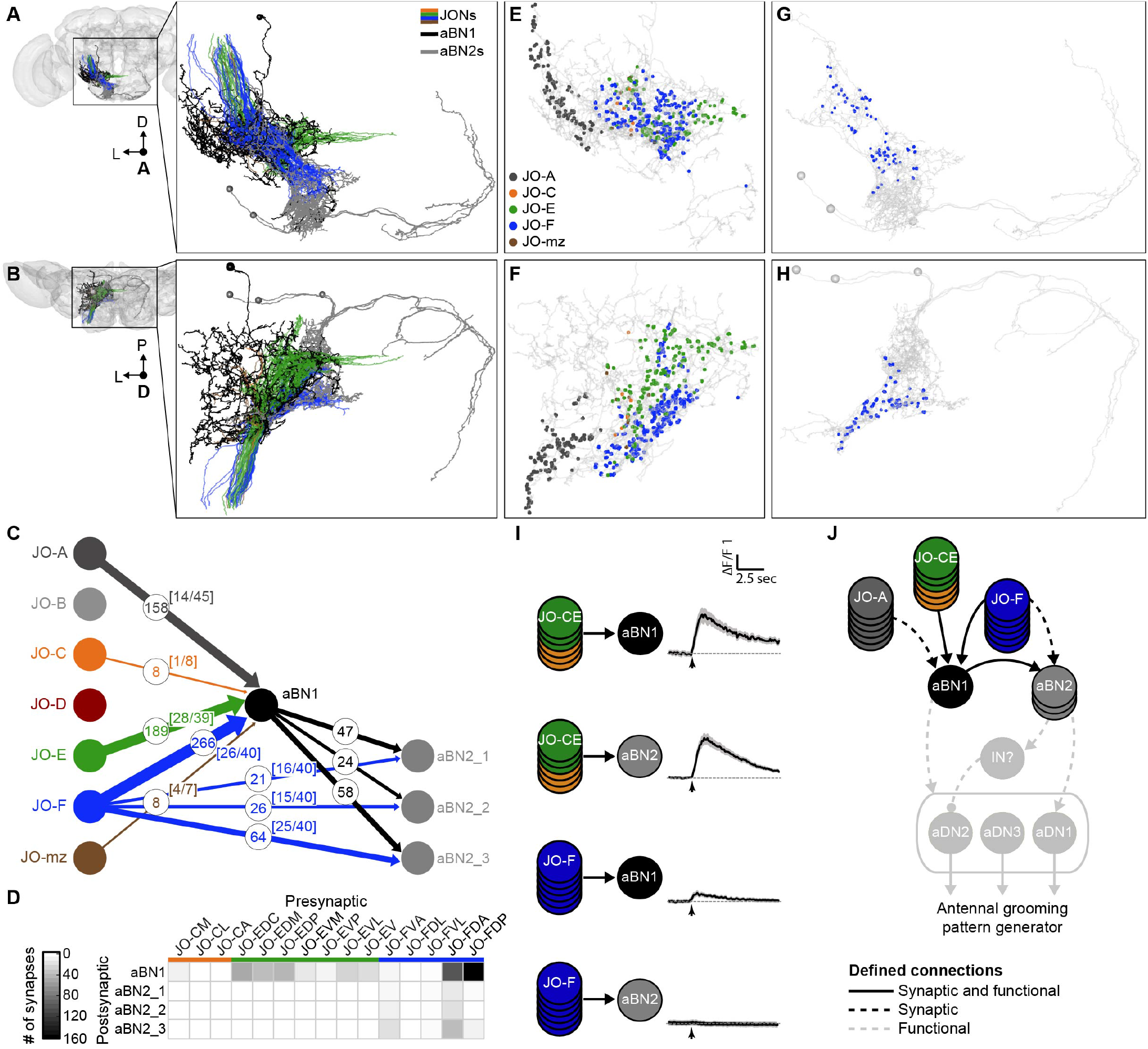
JO-CE and -F neurons are synaptically and functionally connected to aBN1. (**A,B**) Shown are the fully reconstructed JO-C, -E, and -F neurons that are synaptically connected with aBN1 (black) and/or aBN2 neurons (gray). The JON subpopulations are in the same colors shown in **Figure 1B**. Anterior (**A**) and dorsal (**B**) views of these neurons are shown in the brain (left) and magnified (right). (**C**) Axo-dendritic and axo-axonic synaptic connections from the JONs to aBN1 and aBN2 neurons. Nodes representing the JONs, aBN1, and aBN2 are shown as labeled circles. Arrow thickness represents number of synaptic connections, while numbers in the circles indicate the actual number of connections. The number of connected JONs over the total number of fully reconstructed JONs for each subpopulation is shown in brackets. (**D**) Table of synaptic connections of the different JON types (presynaptic) with aBN1 and aBN2 neurons (postsynaptic). The shade of each box corresponds to the total number of synaptic connections indicated on the left. (**E-H**) Spatial organization of synapses of different JON subpopulations onto aBN1 (**E,F**), or all three aBN2 neurons (**G,H**) shown from anterior (**E,G**) and dorsal (**F,H**) views. Colored dots indicate synapses from JO-A (dark gray), JO-C (orange), JO-E (green), and JO-F (blue) neurons. (**I**) JO-CE or -F neurons expressing CsChrimson were exposed to red light while changes in the fluorescence of aBN1 or aBN2 neurons expressing GCaMP6 were imaged. Each tested neuronal pair is shown with labeled circles. Shown is the ΔF/F average ± S.E.M. (n ≥ 5 flies tested with 15 trials per trace). Arrow below each trace indicates when a red light pulse was delivered. (**J**) Revised wiring diagram of the previously published antennal command circuit (Hampel et al., 2015). Arrows represent cholinergic excitatory connections, while the ball and stick represents an inhibitory connection from a yet-to-be identified inhibitory neuron (IN). Arrow to the gray oval around aDN1, 2, and 3 indicates that aBN1 has a functional connection with aDN, but it is unknown to which aDN(s). Solid black lines represent connections that have been defined by synaptic and functional connections. Black dashed lines indicate synaptic connections where the functional connections have not been tested or were not fully tested. Gray dashed lines indicate functional connections that have not been tested for synaptic connections. Solid gray arrows from aDN1,2, and 3 depict presumed descending commands. Note: JO-mz neuron connections with aBN1 are not shown.

In agreement with our proposal that the JO-CE and -F neurons converge onto the antennal command circuit (**Figure 7A**), all three subpopulations have synaptic connections with aBN1 (**Figure 8C**, **Video 5**). However, only 1 of the 9 JO-C neurons was connected (8 synapses), as compared with 28 out of 39 JO-E neurons (189 synapses). This may suggest that the JO-E neurons are the primary mediators of antennal grooming and the JO-C neurons play a more limited role. 26 out of the 40 completely reconstructed JO-F neurons had synapses with aBN1 (266 synapses). The connectivity of the JO-E and -F neurons with aBN1 included almost all of JON types within each subpopulation (**Figure 8D**, **Figure 8 – figure supplement 2**). All of the different JO-E neuron types had connections with aBN1, while all but one of the JO-F neuron types (JO-FDL) had connections. However, JO-FDA and -FDP neurons accounted for 98% of all JO-F-to-aBN1 connections. Thus, multiple different JON types from the subpopulations that elicit antennal grooming have converging synaptic connections onto aBN1.

One surprising result of this analysis was that JONs that were not previously implicated in grooming are connected with aBN1. While the JO-B and -D neurons showed no synaptic connections, 14 out of the 45 reconstructed JO-A neurons (158 synapses) and 4 out of 7 JO-mz neurons (8 synapses) had connections with aBN1 (**Figure 8C**). It remains unclear what role the JO-A and -mz neurons may have in grooming behavior. Our previous experiments suggested that activation of JO-A neurons does not elicit antennal grooming (Hampel et al., 2015). However, it is possible that the driver lines that we used in that study did not express specifically in the JO-A neurons that are connected with aBN1. Future work will be required to test the extent to which the JO-A and -mz neurons have functional inputs onto the antennal command circuit.

aBN1 has a large dendritic field that spans the projection zones of the different JON subpopulations. Thus, because the JONs are topographically organized (**Figure 1B,H**), their synapses onto the aBN1 dendrite reflect this organization (**Figure 8E,F**). Despite this spatial segregation of the JON synapses, both the JO-E and -F neurons synapse onto the same dendritic branches (**Figure 8 – figure supplement 3A**). However, the distribution of their synapses on these branches differed. The JO-F synapses were mainly on the proximal parts of the aBN1 dendrites, while the JO-E synapse were on the distal ends (**Figure 8 – figure supplement 3A,B**). In contrast to the JO-E and -F synapses that were distributed across multiple different aBN1 dendritic branches, synapses from JO-A neurons had a more restricted distribution in the dendritic field (**Figure 8E,F**, **Figure 8 – figure supplement 3A**). This difference in distribution of synapses between the different JON subpopulations could have important implications for how their inputs excite aBN1 or are integrated in the aBN1 dendrite. Previous work has shown that integration at the dendrite has important roles in neural circuit function (Häusser et al., 2000; Magee, 2000).

In contrast to aBN1 that was synaptically connected with most of the different JON subpopulations, only the JO-F neurons had connections with the aBN2 neurons (**Figure 8C,G,H**). However, the JO-F neurons had fewer synapses with the aBN2 neurons than aBN1 (111 synapses with aBN2 neurons versus 266 with aBN1). All three aBN2 neurons received synapses from the JO-F neurons, with aBN2_3 having the highest number (64 synapses), as compared with aBN2_1 and _2 that had 21 and 26 synapses, respectively. All of the JO-F neuron types except the JO-FDL neurons synapsed with the aBN2 neurons (**Figure 8 – figure supplement 2C**). The JO-FDA and -FVA neurons had the majority of synapses, accounting for 86.5% of the JO-F connections with aBN2 neurons. These results suggested that the JO-F neurons could directly influence the antennal command circuit through both aBN1 and aBN2.

The JO-CE and -F neurons all converge onto the antennal command circuit through synaptic connections, however it remained unclear whether these connections were functional. We previously found that the simultaneous optogenetic activation of JO-C, -E, and -F neurons could stimulate calcium responses in aBN1 and aBN2 neurons (Hampel et al., 2015). This demonstrated that at least one of these JON subpopulations was functionally connected with the circuit, however we did not resolve which specific subpopulation(s) were connected. Here, we addressed this question using our previously established functional connectivity assay (Franconville et al., 2018; Hampel et al., 2015). In brief, we measured the calcium responses of aBN1 and aBN2 neurons when the JO-CE or -F neurons were optogenetically activated. This was done using isolated CNSs, where the JON subpopulations expressed CsChrimson (Klapoetke et al., 2014), and aBN1 and aBN2 expressed the calcium responder GCaMP6 (Chen et al., 2013).

Optogenetic activation of the JO-CE neurons evoked calcium responses in both aBN1 and aBN2 neurons (**Figure 8I**, **Figure 8 – figure supplement 4A-E**). This indicated that the aBN1 and aBN2 neurons are functionally connected with the JO-CE neurons. However, the JO-CE neurons are not synaptically connected with aBN2 neurons. This suggests that the functional connectivity is mediated indirectly through the excitation of aBN1 that in turn excites aBN2 neurons (**Figure 8J**). This is supported by our previous work demonstrating that aBN1 is functionally connected with aBN2 neurons (Hampel et al., 2015), and work presented here showing synaptic connectivity between these neurons (129 synapses total, **Figure 8C**).

Optogenetic activation of the JO-F neurons evoked a calcium response in aBN1, while aBN2 neurons showed no response (**Figure 8I**, **Figure 8 – figure supplement 4F,G**). The apparent lack of functional connectivity between the JO-F and aBN2 neurons might be surprising given their synaptic connectivity (**Figure 8C**). The experiment shown in **Figure 8I** was done using JO-F-1 (spGAL4 line) to drive CsChrimson expression in the JO-F neurons, while GCaMP6s was expressed in aBN2 neurons using the published aBN2-LexA driver line (Hampel et al., 2015). We also tested for JO-F to aBN2 functional connectivity using JO-F-3 (LexA line) to drive expression of CsChrimson in the JO-F neurons while GCaMP6s was expressed in the aBN2 neurons using the previously published aBN2-spGAL4-1 driver line (Hampel et al., 2015). Again, we did not observe functional connectivity of the JO-F neurons with aBN2 (not shown). This lack of functional connectivity might be explained as a consequence of the relatively low numbers of synaptic connections between individual JO-F and aBN2 neurons compared with aBN1. The JO-F neurons had an average of 10.2 synapses with aBN1 per neuron (median of 5.5 synapses), and an average of 1.3, 1.7, and 2.7 synaptic connections with aBN2_1, aBN2_2, and aBN2_3 neurons, respectively (median of 1, 1, and 2 synapses). It is unclear if these relatively lower numbers of synapses between the JO-F and aBN2 neurons can account for the lack of observed functional connectivity. However, from this work we can conclude that the JO-CE and -F neurons are both synaptically and functionally connected with aBN1.

### A revised connectivity map of the antennal command circuit

In this study, we provide a neural circuit connectivity-based answer to the question of how distinct subpopulations of JONs could elicit grooming through the antennal command circuit. Based on our previously published calcium imaging studies, we presented a connectivity map of the circuit whereby the JONs were functionally connected with aBN1, aBN2, and at least one of three aDN neurons (Hampel et al., 2015). In the present study, we proposed that aBN1 and aBN2 neurons were the likely synaptic partners of the JONs. We found that the JO-CE and -F neurons converge onto the circuit through synaptic and functional connections with aBN1. However, we also identified synaptic connections from the JO-F neurons to the aBN2 neurons, raising the possibility that this connectivity is partially responsible for mediating the grooming response to JO-F neuron activation. Our reconstructions also revealed that five of the seven JON subpopulations converge onto the circuit (JO-A, -C, -E, -F, and -mz neurons), including two that were not previously implicated in grooming. These new results now enable us to present a revised connectivity map that is based on both synaptic and functional connections and shows how the JONs interface with the antennal command circuit (**Figure 8J**). While our work here identifies chemical synaptic connections, it is possible the JONs also have electrical synapses with the antennal command circuit, which cannot be detected in the EM dataset that we analyzed (Zheng et al., 2018). Indeed, previous studies found that the JO-A neurons have both chemical and electrical synapses with the giant fiber (GF) neurons (Pézier et al., 2014; Strausfeld and Bassemir, 1983). Thus, future work will be required to determine if electrical synapses are also present at the JON-antennal command circuit interface.

The reconstruction of aBN1 and aBN2 neurons has also enabled us to partially define the organization of the early layers of the antennal command circuit. Our previous work had indicated a strong functional connection from aBN1 to the aBN2 neurons, and a relatively weak and inconsistent reciprocal connection from the aBN2 neurons to aBN1. In agreement with this, the synaptic connectivity between these interneurons strongly favored aBN1 to aBN2 connections. That is, there were 134 aBN1 to aBN2 connections, while aBN2_1 and aBN2_3 only had 2 connections each with aBN1 (**Figure 8C**, **Supplementary file 2**). This indicates that the direction of excitation in the circuit is almost exclusively from aBN1 to the aBN2 neurons (**Figure 8J**).

The work presented here does not elucidate the connectivity of the command circuit descending neurons, the aDNs. Our previous anatomical and calcium imaging studies revealed that the three different aDN neurons likely differ in their connectivity within the circuit. First, the dendrites of at least two of the aDNs (aDN1 and aDN2) had different morphologies and resided in different regions of the SEZ. This suggests that the two neurons have different presynaptic partners. Second, calcium imaging revealed that the aDNs differed in their responses to optogenetic activation of upstream neurons. For example, activation of aBN2 caused excitation of aDN1, but induced inhibition in aDN2. The latter result suggested that there was a yet-to-be identified inhibitory neuron in the circuit that was postsynaptic to aBN2 and presynaptic to aDN2 (**Figure 8J**). In addition to revealing likely differential connectivity of the aDNs in the circuit, this result also indicated that at least some of the aDNs are postsynaptic to aBN2 neurons. However, it is possible that the aDNs are synaptically connected with the JONs. For example, the aDN1 dendrites extend dorsally to where the JONs project, and thus could receive direct synaptic connections from JONs. Further, JO-C, -E, and -F activation can induce calcium responses in aDN1, however this connection could be indirect. The EM reconstructions established here will provide the foundation for studies that will address the connectivity of the aDNs within the antennal command circuit.

### The antennal command circuit receives synaptic inputs from neurons other than JONs

Analysis of the dendrites of the reconstructed aBN1 and aBN2 neurons revealed that their postsynaptic sites were not completely occupied by JONs. Only 12% of the postsynaptic sites on aBN1 were occupied by JONs. Given that we have only reconstructed a portion of the JONs, it is likely that some of the unaccounted-for synapses will be from other JONs. Estimates of the number of JO-C and -E neurons from stochastic labeling studies (Kamikouchi et al., 2006) suggest that we reconstructed 47% of the JO-CE neurons (48 out of an estimated 102). However, counts of JONs that are targeted by specific driver lines indicate that there are about 200 JO-CE neurons rather than 102 (Kamikouchi et al., 2006). Based on these different estimates, and assuming that the JO-A and -F neurons are reconstructed to comparable percentages, the postsynaptic sites on aBN1 that are occupied by JONs are between 24 and 50%. Therefore, it is likely that at least 50% of the aBN1 inputs are from neurons that are not JONs. The aBN2 neurons showed an even greater portion of postsynaptic sites that were not occupied by inputs from either aBN1 or JO-F neurons. That is, aBN1 and the JO-F neurons only occupy 3.3%, 1.9%, 5.9% or the postsynaptic sites on the aBN2_1, aBN2_2, and aBN2_3 dendrites, respectively. Although aBN1 occupies a relatively small portion of the aBN2 postsynaptic sites, aBN1 activation is sufficient to elicit calcium responses in aBN2 (Hampel et al., 2015). Further, both aBN1 and aBN2 are necessary for JON-elicited grooming responses. Taken together, our data suggest that we have reconstructed critical neurons in the antennal command circuit, but there are other neurons associated with the circuit that remain to be identified.

What are the functions of these other neurons with inputs into the circuit? One possibility is that mechanosensory neuron types other than the JONs are connected with the circuit to stimulate grooming. The second segment of the antennae has mechanosensory bristles, whose associated mechanosensory neurons project into the same area of the brain as the aBN1 and aBN2 neurons (Homberg et al., 1989; Melzig et al., 1996). Given that stimulating mechanosensory bristles on other body parts elicits site directed grooming responses (Corfas and Dudai, 1989; Vandervorst and Ghysen, 1980), it is possible that the antennal bristle mechanosensory neurons connect with aBN1 and/or aBN2 to elicit grooming. Another possibility is that the unidentified inputs into the circuit are regulatory. We have previously shown that when flies are coated in dust, they groom different parts of their heads and posterior bodies in a particular order (eyes > antennae > abdomen > wings > thorax) (Seeds et al., 2014). This sequence occurs based on a hierarchical suppression mechanism whereby movements that occur earlier in the sequence suppress later ones. One mechanism that we proposed that could drive hierarchical suppression was through inhibitory connections onto the circuits that drive each movement. Thus, future reconstructions will examine the “orphan” postsynaptic sites on aBN1 and aBN2 to identify candidate inhibitory inputs, bristle mechanosensory neurons, and likely other neuronal types.

### JON axons make synaptic connections with each other

Not only did the EM reconstructions in this work reveal how the JON subpopulations connect with the antennal command circuit, but they also revealed that the JONs are synaptically connected with each other. This is not a new observation, as previous work showed that the JONs have axo-axonal electrical and chemical synaptic connections with each other (Sivan-Loukianova and Eberl, 2005). However, analysis of all-to-all connectivity among the different reconstructed JONs revealed that the connectivity only occurred among JONs that belonged to the same subpopulation (**Figure 8 – figure supplement 1**). For example, the JO-F neurons had numerous connections with each other, but showed virtually no connectivity with JO-C or -E neurons. Furthermore, JONs of the same type within a subpopulation were more highly connected with each other than with other types in that subpopulation (e.g. JO-EVP neurons, **Figure 8 – figure supplement 1**). Preferential connectivity of JONs within a particular subpopulation has also been shown with the JO-A and -B neurons (Kim et al., 2020). This type of connectivity among sensory neurons is an emerging theme that is increasingly being described in sensory neurons across modalities (Horne et al., 2018; Marin et al., 2020; Miroschnikow et al., 2018; Tobin et al., 2017). However, the functional significance of this type of connectivity remains unclear.

### Diverse JON subpopulations converge onto the antennal command circuit

Why do different JON subpopulations converge onto the antennal command circuit to elicit grooming? One possible explanation is based on evidence that the subpopulations are tuned to different mechanical stimuli (Ishikawa et al., 2017; Kamikouchi et al., 2009; Mamiya and Dickinson, 2015; Matsuo et al., 2014; Patella and Wilson, 2018). This could provide a means by which diverse stimuli could elicit grooming through the JONs. Different stimuli are known to elicit antennal grooming, including debris on the body surface (e.g. dust) and mechanical displacements of the antennae (Hampel et al., 2015; Phillis et al., 1993; Seeds et al., 2014). Interestingly, the JO-CE neurons are implicated in the grooming response to dust, while the JO-F neurons are implicated in the response to antennal displacements. A recent study showed that expression of tetanus toxin (TNT) using a driver line that appears to target JO-CE neurons disrupted dust induced grooming (Zhang et al., 2020). We found that expression of TNT using a driver that mostly targets the JO-F neurons (aJO-spGAL4-1) disrupted the grooming response to displacements of the antennae (Hampel et al., 2015). However, we could not find evidence that the JO-F neurons respond to such displacements (**Figure 5C,D**). Thus, further work is needed to test the possibility that the JO-CE neurons can detect dust on the antennae, and that the JO-F neurons respond to antennal displacements. Additionally, the JO-A and - mz neurons are connected with the circuit (**Figure 8C**), and the JO-A neurons are known to respond to antennal vibrations (Ishikawa et al., 2017; Kamikouchi et al., 2009; Yorozu et al., 2009). This may suggest that antennal vibrations could also excite the command circuit and influence grooming responses.

### JON involvement in multiple different behaviors

Each JON subpopulation is thought to influence multiple different behaviors. For example, the JO-E neurons are implicated in the suppression of locomotion in response to a wind stimulus (Yorozu et al., 2009), modulating wing movements during flight (Mamiya and Dickinson, 2015), and grooming (**Figure 6A-D**). This raises the question of how the JON types within a subpopulation (e.g. JO-EDC or -EVP neurons) control different behaviors. One possibility is that each type connects with a specific circuit to control a particular behavior (e.g. grooming or flight circuitry). Evidence from the reconstructed JO-A neurons may suggest that this is the case. A subset of JO-A neurons (type-1) interfaces with the jump escape circuitry through connections with the GF neuron (Kim et al., 2020), while the morphologically distinct JO-A type 2 neurons connect with aBN1 in the antennal command circuit (**Supplementary file 2**). In contrast, most of the different JO-E and -F neuron types are connected with aBN1 (**Figure 8 – figure supplement 2C,D**). This may suggest that specific types are not necessarily dedicated to specific behavioral circuits, but are rather connected with multiple different circuits. Our preliminary observations find that not all JON postsynaptic profiles belong to aBN1, aBN2, or JON neurons, revealing that we did not identify all partners of JON presynaptic sites (not shown). Thus, these synapses could connect to unidentified neurons that control other behaviors. However, a closer look at the connectivity between the different JON types and aBN1 may suggest rather that the interface between circuits is governed by a gradient of connection strengths between JON types. For example, all 5 of the completely reconstructed JO-EDC neurons were connected with aBN1, while 4 out of 6 JO-EVP neurons were connected. Further, the 5 JO-EDC neurons have 49 connections onto aBN1 (average of 9.8 synapses per neuron), while the 4 JO-EVP neurons have 6 connections (1.5 synapses). Thus, the extent to which particular JON types connect with multiple different circuits or are dedicated to specific circuits remains an outstanding question.

### A resource for understanding how sensory topography interfaces with neural circuits to influence behavior

The JO is a chordotonal organ in the antennae, but there are also chordotonal organs in other body parts of insects and crustaceans (Field and Matheson, 1998). Chordotonal organs can detect movements of particular limbs for diverse purposes, such as proprioception and sound detection. These mechanosensory structures are studied to address fundamental questions about how stimuli are processed and influence appropriate behavioral responses (Tuthill and Wilson, 2016). There are commonalities among chordotonal organs, as exemplified by recent studies of the fruit fly JO and leg femoral chordotonal organ (FeCO). First, subpopulations of mechanosensory neurons within these chordotonal organs are tuned to specific stimuli, such as vibrations and tonic displacements (Kamikouchi et al., 2009; Mamiya et al., 2018; Patella and Wilson, 2018; Yorozu et al., 2009). Second, these mechanosensory neurons are morphologically diverse and have topographically organized projections into the CNS (Kamikouchi et al., 2006; Mamiya et al., 2018). Third, the subpopulations can differentially interface with downstream circuitry to influence distinct behaviors or movements (Agrawal et al., 2020; Hampel et al., 2015; Kim et al., 2020; Vaughan et al., 2014). Fourth, similar features of mechanosensory stimuli can be represented in neurons downstream of the JO and FeCO (Agrawal et al., 2020; Chang et al., 2016). These commonalities suggest that results obtained through studies of different chordotonal organs could be mutually informative. However, there is a dearth of information about how chordotonal mechanosensory neurons interface with downstream circuitry at the synaptic level. Our work here, along with two previous studies (Kim et al., 2020; Maniates-Selvin et al., 2020) reveal the near complete topography of mechanosensory neurons that make up the JO and the FeCO. This provides the foundation for rapid identification of neural circuitry that is postsynaptic to two different chordotonal organs. A synaptic resolution view of the interface between the JO and FeCO and downstream circuitry will provide a valuable resource for addressing fundamental questions about the functional significance of mechanosensory topography.

## Materials and methods

### Rearing conditions and fly stocks

The GAL4, spGAL4, and LexA lines that were used in this study were generated by the labs of Gerald Rubin and Barry Dickson and most lines can be obtained from the Bloomington *Drosophila* stock center (Dionne et al., 2017; Jenett et al., 2012; Pfeiffer et al., 2008; Tirián and Dickson, 2017). Stocks carrying multiple transgenes that were built in this study are available on request. Control flies contain the DNA elements used for generating the different spGAL4 halves or LexA collections, but lack enhancers to drive their expression (Pfeiffer et al., 2008, 2010). The complete list of fly stocks that were used in this study can be found in the Key resources table.

GAL4, spGAL4, and LexA lines were crossed to their respective UAS or LexAop driver lines. Flies were reared on cornmeal and molasses food at 21 °C and 50-60% relative humidity on a 16/8-hour light/dark cycle. Flies that were used for optogenetic experiments were reared on food containing 0.4 mM all-*trans*-retinal in vials that were wrapped in aluminum foil and covered with a box to keep them in the dark. Unless otherwise stated, flies used for experiments were male and 5 to 8 days old.

### Neural circuit reconstructions from an EM volume

Neurons and their synapses were reconstructed from a serial section transmission electron microscopy volume of a female full adult fly brain (FAFB) at 4 x 4 x 40 nm resolution (Zheng et al., 2018). All reconstructions were done by an experienced tracer who used two different approaches. The first approach was based on manual annotation and provides complete reconstruction of the neurites and pre- and postsynaptic sites of each neuron. The browser-based software CATMAID (http://catmaid.org) (Saalfeld et al., 2009) was used to manually navigate through the volume image stacks and manually place nodes that marked the neurites and synapses (Schneider-Mizell et al., 2016). For synapse annotations, we followed the criteria used by the FAFB connectomics community. Briefly, synapses had to show at least three out of the four following features: 1) an active zone with presynaptic vesicles, 2) a clear presynaptic density (such as a ribbon or T-bar), 3) a synaptic cleft, and 4) a postsynaptic density. For further tracing guidelines see Zheng et al., 2018. Manual tracing had the disadvantage of being labor intensive, which limited the number of neurons that could be reconstructed. Therefore, we employed the second approach of using an automated segmentation algorithm that uses flood-filling networks (Li et al., 2019). The algorithm would occasionally create false splits. Therefore, the tracer resolved these false splits by manually assembling the fragments as previously described (Marin et al., 2020). This approach enabled us to semi-automatically annotate the major branches of each neuron, but not the fine branches and synaptic sites.

To locate the JON subpopulations in the EM volume, we first registered a light-microscopy confocal z-stack of these neurons into the volume. The z-stack was of a transgenic driver line (R27H08-GAL4) that expresses in the JO-C, -E, and -F neurons (Hampel et al., 2015). R27H08-GAL4 (RRID: BDSC_49441) was crossed to *10XUAS-IVS-mCD8::GFP* (RRID: BDSC_32185) to label these JONs with GFP. The brains were dissected, stained, and imaged by confocal microscopy as described below (**Figure 1 – figure supplement 1A**). The resulting image stack was registered into the EM volume using the software ELM (Bogovic et al., 2016, 2018) to highlight the JON axons where the antennal nerve enters the brain as a neuron bundle. The medial region of the bundle, where JO-C and -E neurons were previously described to project (Kamikouchi et al., 2006), was highlighted by GFP (**Figure 1 – figure supplement 1B**).

We next reconstructed 147 JONs within the GFP-highlighted region (**Figure 1A,B**, **Figure 1 – figure supplement 1C**, colored dots). Neuron reconstructions were performed until we had identified JO-C, -D, -E, and -F neurons, and could not uncover new morphologically distinct JONs with further reconstructions. 70 JONs were manually reconstructed to completion, including all of their pre- and postsynaptic sites. We then reconstructed 77 additional JONs by assembling fragments created by the automated segmentation algorithm. 34 of these JONs were proofread using previously published methods (Schneider-Mizell et al., 2016), and traced to completion. Thus, out of the 147 reconstructed JONs, 104 (71%) were completely reconstructed with their entire morphology and all pre- and postsynaptic sites. At least 63% of the reconstructed JONs for each subpopulation were fully reconstructed (**Figure 1 – figure supplement 2-6**, neurons marked with asterisks). Reconstructed JONs were next categorized by manual annotation and named. Manual annotations were done by comparing the morphology and projections of the reconstructed JONs with published light microscopy studies (Hampel et al., 2015; Kamikouchi et al., 2006). We categorized the reconstructed JONs into 17 different types (140 JONs), and a group of 7 neurons innervating multiple zones. We estimate that, of the 173 previously published numbers of JO-C, -D, -E, and mz neurons (Kamikouchi et al., 2006), we have reconstructed 50% (37% fully reconstructed). See **Supplementary file 1** for detailed information on each JON type, including their FAFB skeleton ID numbers, raw and smooth cable length, number of nodes, and number of pre- and postsynaptic sites.

We next performed an NBLAST all-to-all comparison of the 147 reconstructed JONs (Costa et al., 2016). We first pruned the primary axonal branch of each JON from its start point in the antennal nerve to its first branch point. Next, twigs shorter than 1 μm were pruned (**Figure 2B**, pruned neurons shown in black in the first panel). The pruning enabled us to cluster the synapse rich parts of the JONs while adjusting for any differences in neuron morphology between the manual reconstruction and automated segmentation methods. At a cut height of h = 1.4, NBLAST clustered the JONs into 15 groups that were mostly consistent with the JON types that we had identified by manual annotation (**Figure 2A,B**).

To identify the brain neurons (i.e. aBN1 and aBN2) in the antennal command circuit (Hampel et al., 2015), we traced neuron profiles found postsynaptic to specific JONs. We identified and completely manually reconstructed a candidate aBN1 neuron postsynaptic to a JO-E neuron, JO_EDM_07 (**Figure 1 – figure supplement 4B**). This candidate neuron was confirmed to be aBN1 by comparing its morphology with confocal images of aBN1 from our previous work (Hampel et al., 2015). Our searches in the EM volume only identified one aBN1 neuron. Our limited searches of postsynaptic partners of different JONs did not identify any aBN2 neurons. This led us to search for aBN2 postsynaptic to aBN1. We identified a neuron that was morphologically consistent with the MCFO-labeled aBN2s postsynaptic to aBN1 (named aBN2_1, **Figure 7F**). This neuron was completely manually reconstructed. Our previous work indicated that the aBN2 axons project through the brain as a bundle. Therefore, we searched around the axon of reconstructed aBN2_1 and identified two morphologically similar neurons that we named aBN2_2 and aBN2_3 (**Figure 7G,H**). These two neurons were initially partially reconstructed by assembling fragments created by the automated segmentation algorithm. The neurons were then proofread and manually traced to completion.

Neurons were plotted and their connectivity analyzed using the natverse package (http://natverse.org/) (Bates et al., 2020) in R version 3.6.2. For the analysis of the JON, aBN1, and aBN2 connectivity, we excluded dendro-axonic and dendro-dendritic connections. This was done by performing an axon-dendrite-split of aBN1 and the aBN2s using synapse flow centrality (Schneider-Mizell et al., 2016) in CATMAID. We then analyzed the locations of pre- and postsynaptic sites along the neuron arbor to establish the type of connections with other neurons. This was done by filtering out all presynapses on the dendrite. For analysis of connectivity with the JONs, the entirety of JON projections into the brain were considered to be axons. Dendrograms of the JON connections with aBN1 were plotted using code by Markus Pleijzier (https://github.com/markuspleijzier/AdultEM/tree/master/Dendrogram_code) (Felsenberg et al., 2018) with the pymaid package (https://github.com/schlegelp/pymaid) in Python version 3.7.4. For visualization of the AMMC neuropile, an alpha-shape was created from all nodes of the reconstructed mechanosensory neurons (147 JON skeletons in this study and the 90 JO-A and -B skeletons from (Kim et al., 2020)) and transformed into a mesh object in R (alphashape3d and rgl packages). The neurons were rendered for videos in Blender version 2.79 with the CATMAID-to-Blender plugin (https://github.com/schlegelp/CATMAID-to-Blender) (Schlegel et al., 2016).

### Identification of driver lines that express in JON subpopulations that elicit antennal grooming

We used a Split GAL4 (spGAL4) screening approach to produce driver lines that expressed in JO-C, - E, and -F neurons. The spGAL4 system enables independent expression of the GAL4 DNA binding domain (DBD) and activation domain (AD). These domains can be reconstituted into a transcriptionally active protein when they are expressed in the overlapping cells of two different patterns (Luan et al., 2006; Pfeiffer et al., 2010). We expressed the DBD in JO-C, -E, and -F neurons using the R27H08 enhancer fragment (R27H08-DBD, RRID: BDSC_69106). To target specific subpopulations of JONs within this pattern, we identified candidate lines that were predicted to express the AD in JO-C, -E, or -F neurons (Dionne et al., 2017; Tirián and Dickson, 2017). This was done by visually screening through a database of images of the CNS expression patterns of enhancer-driven lines (Jenett et al., 2012). About 30 different identified candidate-ADs were crossed to flies carrying R27H08-DBD and *20xUAS-CsChrimson-mVenus* (RRID: BDSC_55134) (Klapoetke et al., 2014). The progeny of the different AD, DBD, and *20xUAS-CsChrimson-mVenus* combinations were placed in behavioral chambers and exposed to red light (optogenetic activation methods described below). We tested three flies for each combination, a number that we previously found could identify lines with expression in neurons whose activation elicit grooming. We then stained and imaged the CsChrimson-mVenus expression patterns of the brains and ventral nervous systems of AD/DBD combinations that elicited grooming (immunohistochemistry and imaging methods described below). Four different DBD/AD combinations were identified that expressed in either zone C/E- or F-projecting JONs. The four AD “hits” were 100C03-AD (aka VT005525, RRID: BDSC_72267), R39H04-AD (RRID: BDSC_75734), R25F11-AD (RRID: BDSC_70623), and 122A08-AD (aka VT050231, RRID: BDSC_71886).

We produced lines that contained both the AD and DBD in the same fly. Two of these lines express in JO-C and -E neurons and were named JO-CE-1 (100C03-AD ⋂ R27H08-DBD) and JO-CE-2 (R39H04-AD ⋂ R27H08-DBD) (**Figure 3A,B**). The other two lines express mainly in JO-F neurons and were named JO-F-1 (R25F11-AD ⋂ R27H08-DBD) and JO-F-2 (122A08-AD ⋂ R27H08-DBD) (**Figure 3C,D**). In a different search, we screened through the image database described above to identify a LexA driver line, R60E02-LexA (RRID: BDSC_54905), that expresses specifically in JO-F neurons (named JO-F-3). See Key resources table for more information about these driver line stock sources and references.

### Immunohistochemical analysis of the driver line expression patterns in the CNS and antennae

We evaluated the expression patterns of the different GAL4, spGAL4, and LexA driver lines using the same staining protocol. GFP or Venus-tagged CsChrimson (for spGAL4 driver line screening only) were expressed by crossing the lines to either *10XUAS-IVS-mCD8::GFP, 20xUAS-CsChrimson-mVenus*, or *13xLexAop-myr::GFP* (RRID: BDSC_32209). The brains, VNCs, and antennae were dissected and stained as previously described (Hampel et al., 2011, 2015). The brains and VNCs were stained using anti-GFP and anti-nc82 antibodies, while the antennae were stained using anti-GFP and anti-ELAV. The following primary and secondary antibodies were used for staining: rabbit anti-GFP (1:500, Thermo Fisher Scientific, Waltham, MA, #A11122), mouse mAb anti-nc82 (1:50, Developmental Studies Hybridoma Bank, University of Iowa), mouse anti-ELAV and rat anti-ELAV (used together for the antennal stain, 1:50, Developmental Studies Hybridoma Bank), AlexaFluor-488 (1:500; goat anti-rabbit; Invitrogen, Carlsbad, CA), and AlexaFluor-568 (1:500; goat anti-mouse, goat anti-rat; Invitrogen).

For multicolor flipout (MCFO) experiments, JO-CE-1, JO-CE-2, JO-F-1, JO-F-2, and aBN2-spGAL4-1 (Hampel et al., 2015) were crossed to the MCFO-5 stock (RRID: BDSC_64089) (Nern et al., 2015). 1 to 3-day old fly brains were dissected and stained using anti-V5, -FLAG, and -HA antibodies. The following primary and secondary antibodies were used: rat anti-FLAG (Novus Biologicals, LLC, Littleton, CO, #NBP1-06712), rabbit anti-HA (Cell Signaling Technology, Danvers, MA, #3724S), mouse anti-V5 (Serotec, Kidlington, England #MCA1360), AlexaFluor-488 (1:500; goat anti-rabbit, goat anti-chicken, goat anti-mouse; Invitrogen), AlexaFluor-568 (1:500; goat anti-mouse, goat anti-rat; Invitrogen), AlexaFluor-633 (1:500; goat anti-rat; Invitrogen). We imaged individually labeled neurons from at least 10 brains for each line. Note: we made several attempts to obtain individually labeled JONs that were part of the JO-CE-1 and −2 expression patterns. However, all of the brains that we examined showed labeling of too many neurons to visualize any one JON.

Stained CNSs and antennae were imaged using a Zeiss LSM800 confocal microscope (Carl Zeiss, Oberkochen, Germany). Image preparation and adjustment of brightness and contrast were performed with Fiji software (http://fiii.sc/). For visualizing the imaged JONs together as shown in **Figure 3H,I**, individual confocal stacks of the different spGAL4 lines were computationally aligned to the JFRC-2010 standard brain (www.virtualflybrain.org) using the Computational Morphometry Toolkit (CMTK) (https://www.nitrc.org/projects/cmtk/) (Jefferis et al., 2007). The aligned confocal stacks were then assembled in FluoRender (Wan et al., 2009, 2012), a suite of software tools for viewing image data. We compared the morphology of the imaged JONs, aBN1, and aBN2 neurons that were imaged via confocal microscopy with their corresponding EM reconstructed neurons using FIJI and CATMAID, respectively.

### Testing the responses of JO-CE and JO-F neurons to stimulations of the antennae

We tested the responses of the JON subpopulations to mechanical stimulations of the antennae using a previously published preparation (Matsuo et al., 2014). The JO-CE-1, JO-CE-2, JO-F-1, and JO-F-2 driver lines were crossed to *UAS-GCaMP6f* flies (RRID: BDSC_42747) (Chen et al., 2013). The progeny were cold anesthetized on ice for one minute and then attached to an imaging plate using silicon grease (SH 44M; Torray, Tokyo, Japan) with the dorsal side up. The proboscis was removed to access to the ventral brain for monitoring changes in fluorescence (Yamada et al., 2018). To prevent dehydration of the brain, saline solution was applied to the opening of the head. The solution contained 108 mM NaCl, 5 mM KCl, 2 mM CaCl2, 8.2 mM MgCl2, 4 mM NaHCO3,1 mM NaH2PO4, 5 mM trehalose, 10 mM sucrose, and 5 mM HEPES, and was adjusted to pH 7.5 with 1 M NaOH, and 265 mOsm (Wang et al., 2003). Neural activity was monitored using a fluorescence microscope (Axio Imager.A2; Carl Zeiss, Oberkochen, Germany) equipped with a water-immersion 20x objective lens [W Achroplan/W N-Achroplan, numerical aperture (NA) 0.5; Carl Zeiss], a spinning disc confocal head CSU-W1 (Yokogawa, Tokyo, Japan), and an OBIS 488 LS laser (Coherent Technologies, Santa Clara, CA) with an excitation wavelength of 488 nm as previously described (Yamada et al., 2018).

Antennal displacements were induced using electrostatic forces that were generated using electrodes (Albert et al., 2007; Effertz et al., 2012; Kamikouchi et al., 2009, 2010). The electrical potential of the fly was increased to +15 V against ground via a charging electrode, a 0.03 mm diameter tungsten wire (Nilaco, Tokyo, Japan) that was inserted into the thorax. The following voltage commands were used: (1) sinusoids of various frequencies (200, 400, and 800 Hz), ranging from −14 V to +14 V, and (2) positive and negative steps, −50 V and +50 V for static push and pull deflections. These stimuli were applied for 4 seconds to a stimulus electrode, a 0.3 mm diameter platinum wire (Nilaco, Japan) that was placed in front of the arista of the fruit fly (Matsuo et al., 2014). At the end of the experiment, samples that did not show responses to any of the tested stimuli were treated with 50 μL of 4.76 M KCl that was pipetted into the saline solution (2 mL volume).

Images were acquired at a rate of 10 Hz with a 100 ms exposure time. F_0_ was defined as the F value obtained 2.5 seconds before the stimulus onset. Four trials were run for each stimulus in a single fly and then averaged. 10 or 12 flies were tested for each driver line for the push, pull and 200 Hz sinusoids. 2 flies were tested for the 400 and 800 Hz sinusoids. To compare the responses between “No stimulation” (NoStim) and “Stimulation” (Stim) conditions, we used 40 frames (4 seconds) before the stimulus onset for No stim and 40 frames (4 sec) during the stimulus for Stim. The Wilcoxon signed-rank test was applied for the statistical analysis of the data.

### Behavioral analysis procedures

We tested for behavioral changes that are caused by activating either JO-CE or -F neurons. The JO-CE-1, JO-CE-2, JO-F-1, and JO-F-2 driver lines were crossed to *20xUAS-CsChrimson-mVenus* (attP18, RRID: BDSC_55134) and *13XLexAop2-IVS-CsChrimson-mVenus* (attP18, RRID: BDSC_55137). The optogenetic behavioral rig, camera setup, and methods for the recording and behavioral analysis of freely moving flies were described previously (Hampel et al., 2015; Seeds et al., 2014). In brief, we used 656-nm red light at 27 mW/cm^2^ intensity (Mightex, Toronto, Canada) for activation experiments using CsChrimson. The red-light stimulus parameters were delivered using a NIDAQ board controlled through Labview (National Instruments, Austin, TX). The red-light frequency chosen was 5 Hz for 5 s (0.1 second on/off), and 30 s interstimulus intervals (total of 3 stimulations). Manual scoring of grooming behavior captured in prerecorded video was performed with VCode software (Hagedorn et al., 2008) and the data was analyzed in MATLAB (MathWorks Incorporated, Natick, MA). Antennal grooming was scored as previously described (Hampel et al., 2015; Seeds et al., 2014), however, in this work the wing flapping and avoidance responses are newly described. Avoidance was scored when the fly body moved backwards by any amount. Wing flapping was scored when the wings start moving to the sides, or up and down until there is no further vibrating movement detectable. Behavioral data was analyzed using nonparametric statistical tests. We performed a Kruskal-Wallis (ANOVA) test to compare more than three genotypes with each other. After that we used a post-hoc Mann-Whitney U test and applied Bonferroni correction. The changes in grooming that we observed by activating the different JON subpopulations had a comparable effect size to our previously published work (Hampel et al., 2015). Therefore, at least 10 experimental and 10 control flies were tested (95% power to detect a 1.48 effect size at a 0.05 significance level).

### Functional connectivity between JON subpopulations and aBN1 or aBN2

This experiment required the use of two binary expression systems (GAL4/UAS and LexA/LexAop) for driving the expression of CsChrimson and GCaMP6 in different neurons in the same fly. For the experiment shown in **Figure 8I**, CsChrimson was expressed in the JON subpopulations using LexA and spGAL4 driver lines that were identified in this study that were specific for either JO-CE or -F neurons. LexA and spGAL4 driver lines that were used for expressing GCaMP6 in aBN1 (aBN1-spGAL4-1) and aBN2 (aBN2-LexA) were identified in our previous study (Hampel et al., 2015). The different driver lines that were used have already been introduced above, with the exception of a LexA driver line that expresses in JO-CE neurons. To identify a driver line that expresses in JO-CE neurons, we screened through the image database described above to identify a LexA driver line, 100C03-LexA (Tirián and Dickson, 2017). This line was named JO-CE-3 (**Figure 8 – figure supplement 5A-C**).

The following stocks were used to test for functional connectivity between the different indicated pairs and for the control. Control: spGAL4 control (pBPp65ADZpUw (attP40), pBPZpGAL4DBDUw (attP2)), 26B12-LexA, *20XUAS-CsChrimson-mVenus* (attP18), *13XLexAop2-IVS-p10-GCaMP6s* (su(Hw)attP5) (RRID: BDSC_44589). JO-CE to aBN1: JO-CE-3 (100C03-LexA (attP40)), aBN1-spGAL4-1, *13XLexAop2-CsChrimson-tdTomato* (VK00005) (RRID: BDSC_82183), and *20xUAS-IVS-GCaMP6m* 15.629 (attP2). JO-CE to aBN2: JO-CE-1, aBN2-LexA, *20XUAS-CsChrimson-mVenus* (attP18), *13XLexAop2-IVS-p10-GCaMP6s* (su(Hw)attP5). JO-F to aBN1: JO-F-3, aBN1-spGAL4-1, *13XLexAop2-CsChrimson-tdTomato* (attP18), *20XUAS-IVS-GCaMP6s* (VK0005) (RRID: BDSC_42749). JO-F to aBN2: JO-F-1, aBN2-LexA, *20XUAS-CsChrimson-mVenus* (attP18), *13XLexAop2-IVS-p10-GCaMP6s* (su(Hw)attP5).

Functional connectivity experiments were performed as previously described (Hampel et al., 2015). Regions of interest of imaging GCaMP6 were chosen that contained distinctive process of either aBN1 or aBN2 neurons (**Figure 8 – figure supplement 4A,B**). CsChrimson was excited by delivering 2 ms pulses of 590-nm light through the objective via an LED. Instantaneous powers measured from the objective were either trains at 50 μW/mm^2^ or 500 μW/mm^2^ delivered at 5 Hz. Experiments started with a 20-pulse train at 50 μW/mm^2^. When no response or a weak response was observed, the power was increased to 500 μW/mm^2^. Experimental runs consisted of four repeats lasting for approximately 20 seconds. Runs were repeated approximately every 2 minutes. When postsynaptic responses were observed, we did not see any desensitization. ΔF/F0 was calculated as previously described (Hampel et al., 2015). The analysis was run in Julia (http://julialang.org/).

## Acknowledgements

We thank Gerald Rubin and Barry Dickson for providing split GAL4 lines, Eric Hoopfer for MATLAB code used to analyze behavioral data, Steven Sawtelle for constructing the behavioral rig and optogenetic setup, Karen Hibbard and Todd Laverty for flies and organizational help, Tom Kazimiers for development and maintenance of CATMAID, John Bogovic for aligning light microscopy images to the EM dataset using ELM, Zari Zavala-Ruiz and the Janelia Visiting Scientist Program for enabling our initial circuit reconstructions, Greg Jefferis and Marta Costa for hosting Katharina Eichler at Cambridge University, Maria Sosa and Bethzaida Birriel for help with visitor agreements and visas, Steven Calle-Schuler for videos, Markus Pleijzier for help with the BN dendrograms, Gwyneth Card and the Janelia tracers for contributing reconstructions of JO-D neurons. This work was funded by the Whitehall Foundation (2017-12-69), NIMHD MD007600 (RCMI), and NIH NIGMS GM103642.

---

**Figure 1 – figure supplement 1.**
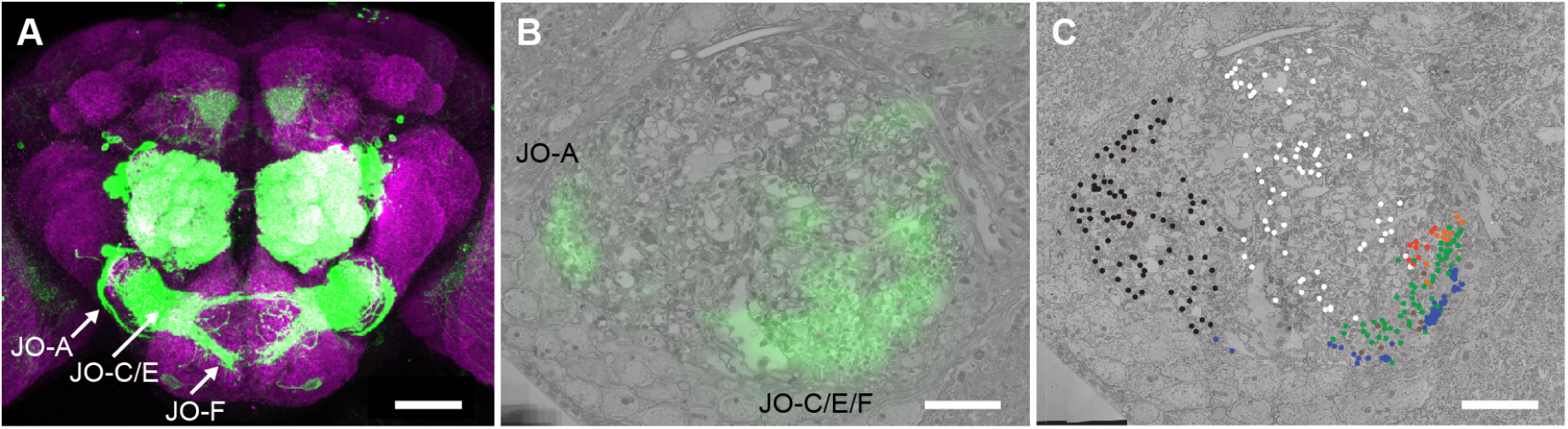
Identifying JONs in the EM volume. (**A**) Brain of R27H08-GAL4 expressing GFP. Shown is the maximum intensity projection of a co-stain with anti-GFP (green) and anti-Bruchpilot (magenta) antibodies to visualize the JON afferent projections into the brain neuropile. Scale bar, 100 μm. (**B**) Registration of R27H08-GAL4 expression pattern into the EM volume to locate JONs implicated in antennal grooming. Shown is an EM section of the point where the antennal nerve enters the brain. The labels indicate the GFP highlighted regions that include either JO-A or -C, -E, and -F neurons. Scale bar, 10 μm. (**C**) Reconstruction of JONs in the medial region highlighted by the R27H08-GAL4 expression pattern. Colored dots indicate individual reconstructed JONs that were identified as JO-A (black), -B (white), -C (orange), -D (red), -E (green), -F (blue) neurons, and JONs that projected to multiple zones (brown). Scale bar, 10 μm.

**Figure 1 – figure supplement 2.**
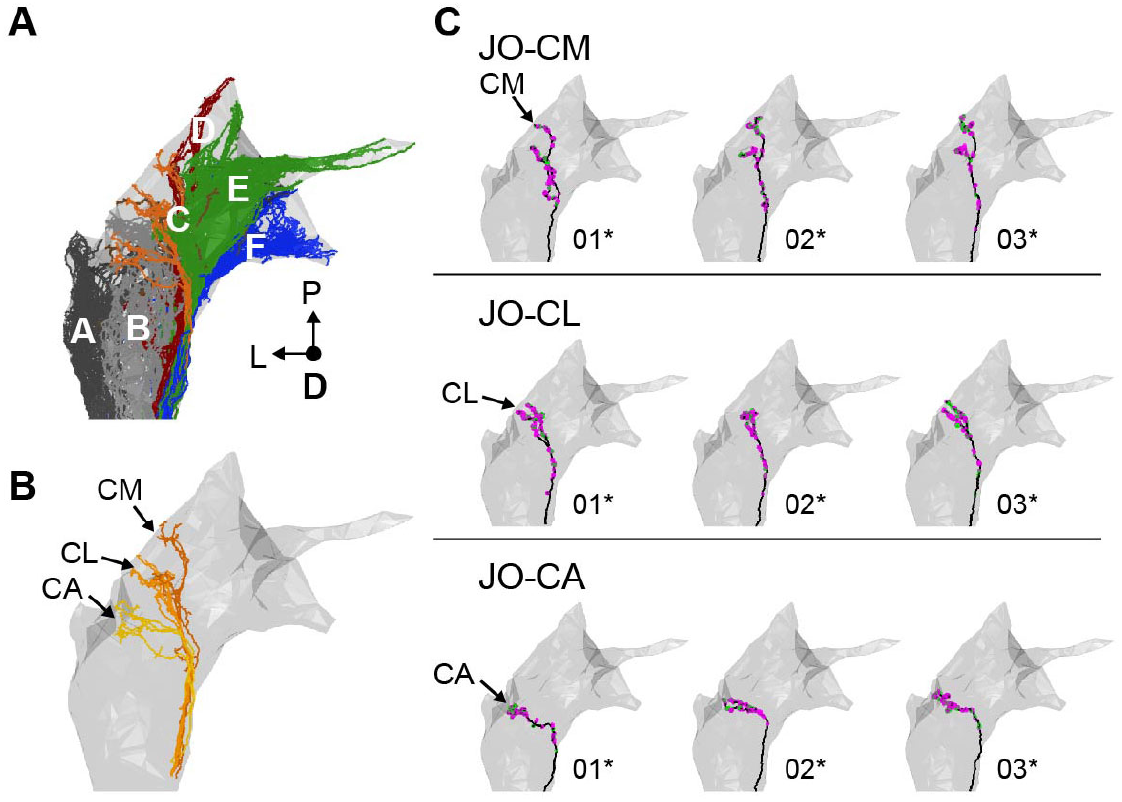
Individual reconstructed zone C-projecting JONs. (**A**) Dorsal view of all reconstructed JONs from the EM dataset with the mesh that outlines the JON neuropile (the gray mesh is used in **B** and **C**, **Figure 1**, and **Figure 1 – figure supplements 3-6**). The colors correspond to the zones to which the different JONs project, including zones A (black), B (gray), C (orange), D (red), E (green), F (blue), and multiple zones (brown). (**B**) Dorsal view of all reconstructed JONs that project to zone C (described in **Figure 1C**). Zone subareas are indicated with labeled arrows. (**C**) Dorsal views of individual zone C-projecting JON types. The JON type is labled for each row. The subarea that receives the larges JON branch is labeled for each type. Numbers below each image indicate the corresponding reconstructed neuron in the EM dataset (e.g. JO_CM_01). Asterisks by the numbers indicate the JONs that have been completely reconstructed, including pre- and postsynaptic sites. Pre- and postsynaptic sites are labeled in magenta and green, respectively.

**Figure 1 – figure supplement 3.**
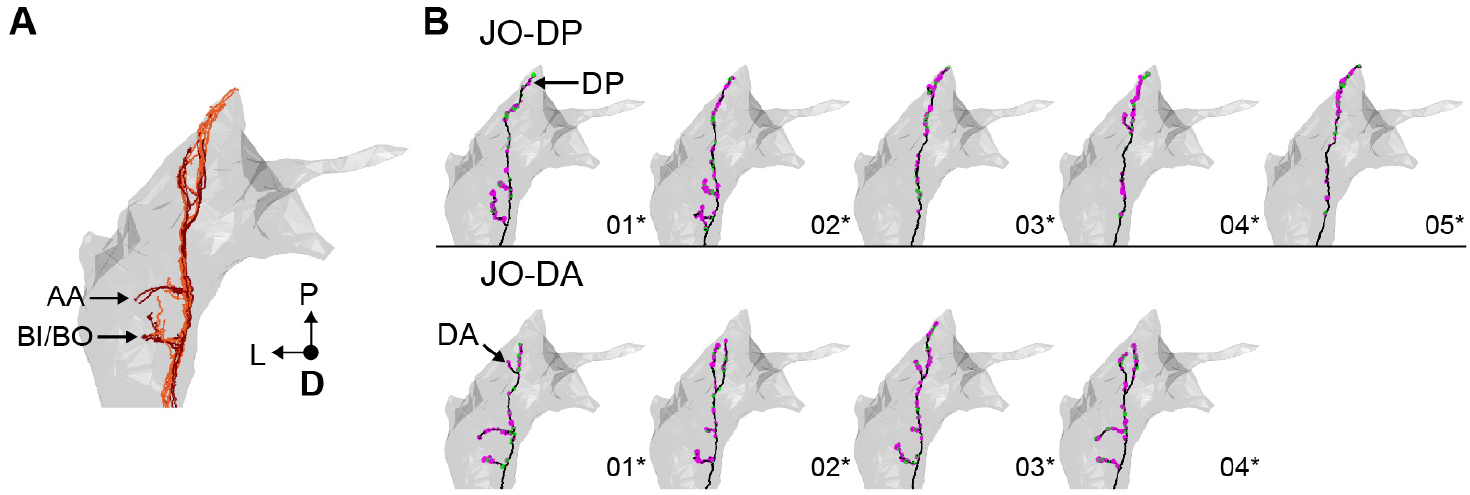
Individual reconstructed zone D-projecting JONs. (**A**) Dorsal view of all reconstructed JONs that project to zone D (described in **Figure 1D**). Zone subareas are indicated with labeled arrows. (**B**) Dorsal views of individual zone D-projecting JON types. The JON type is labled for each row. Numbers below each image indicate the corresponding reconstructed neuron in the EM dataset (e.g. JO_DP_01). Asterisks by the numbers indicate the JONs that have been completely reconstructed, including pre- and postsynaptic sites. Pre- and postsynaptic sites are labeled in magenta and green, respectively.

**Figure 1 – figure supplement 4.**
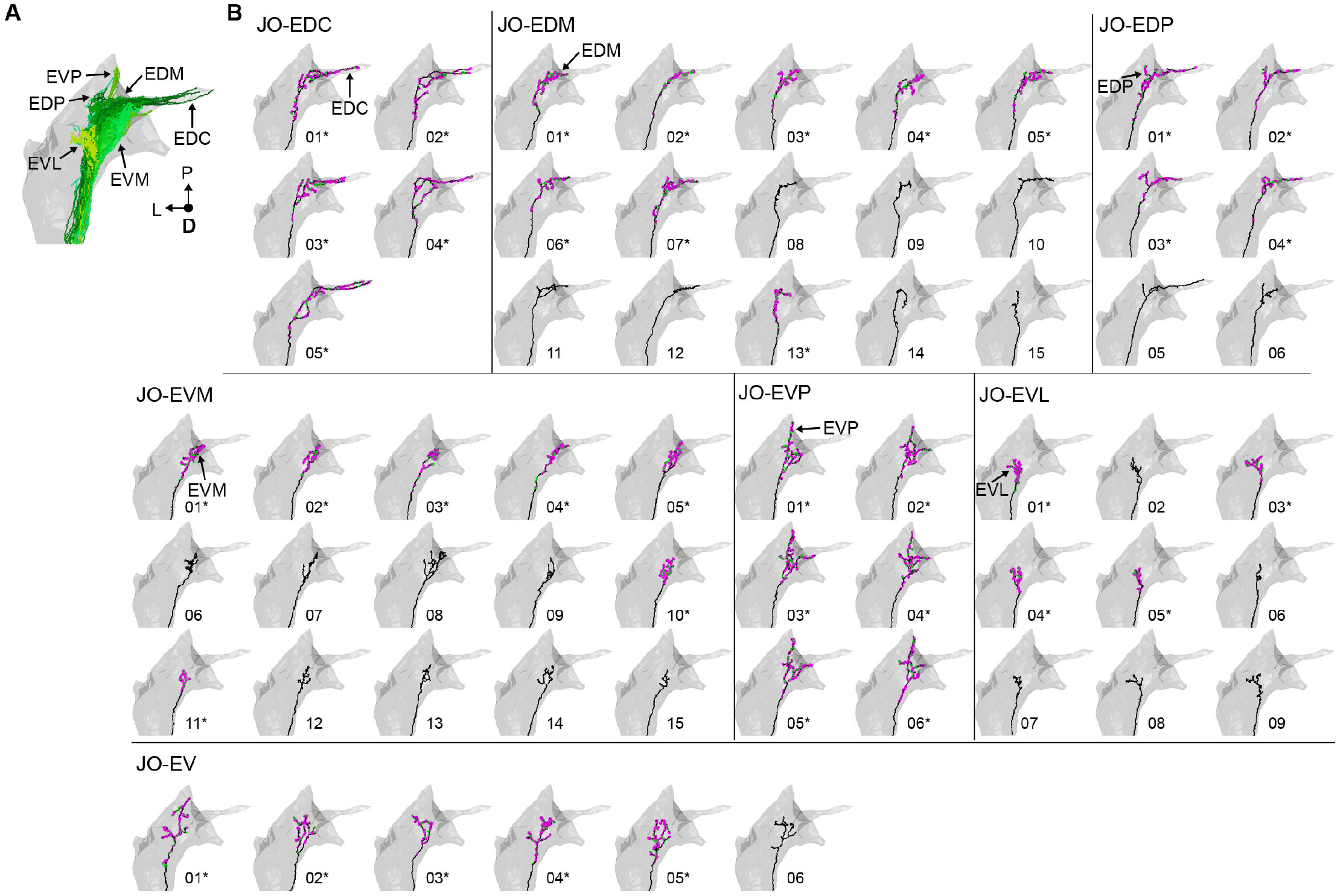
Individual reconstructed zone E-projecting JONs. (**A**) Dorsal view of the reconstructed JONs that project to zone E (described in **Figure 1E**). Zone subareas are indicated with labeled arrows. (**B**) Dorsal view of individual zone E-projecting JON types. Arrows with labels indicate the subarea that the JON projects to that led to the naming of the JON. The names of each of the seven JON types are labeled. Numbers below each image indicate the corresponding reconstructed neuron in the EM dataset (e.g. JO_EVM_01 or JO_EDP_05). Asterisks by the numbers indicate the JONs that have been completely reconstructed, including pre- and postsynaptic sites. Pre- and postsynaptic sites are labeled in magenta and green, respectively.

**Figure 1 – figure supplement 5.**
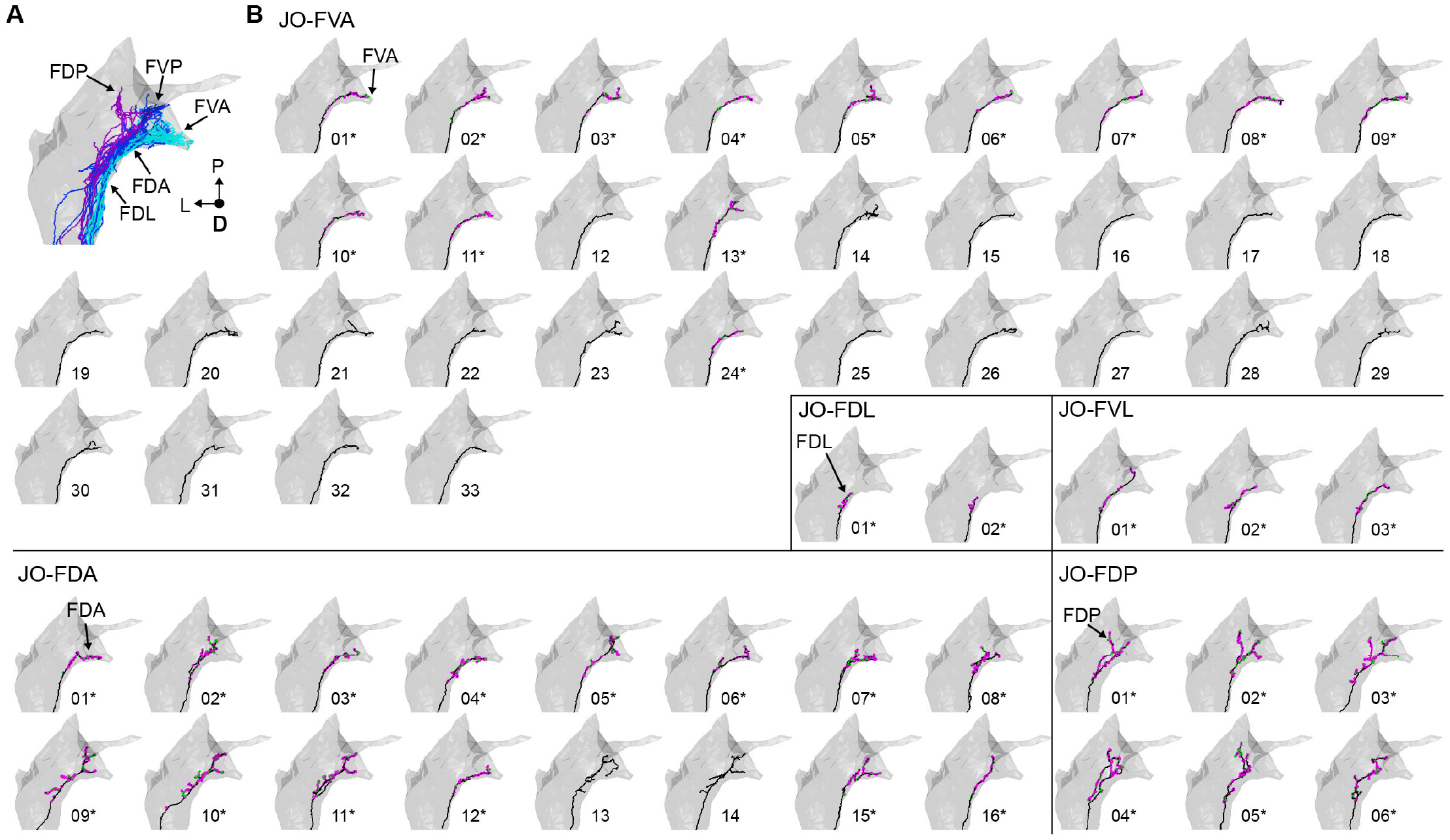
Individual reconstructed zone F-projecting JONs. (**A**) Dorsal view of the reconstructed JONs that project to zone F (described in **Figure 1F**). Zone subareas are indicated with labeled arrows. (**B**) Five different types of zone F-projecting JONs are shown from a dorsal view. The name of each of the five JON types is labeled. Arrows with labels indicate the subarea that the JON projects to that led to the naming of the JON. Numbers below each image indicate the corresponding reconstructed neuron in the EM dataset (e.g. JO_FVA_01 or JO_FDA_02). Asterisks by the numbers indicate which JONs have been completely reconstructed, including pre- and postsynaptic sites. Pre- and postsynaptic sites are labeled in magenta and green, respectively.

**Figure 1 – figure supplement 6.**
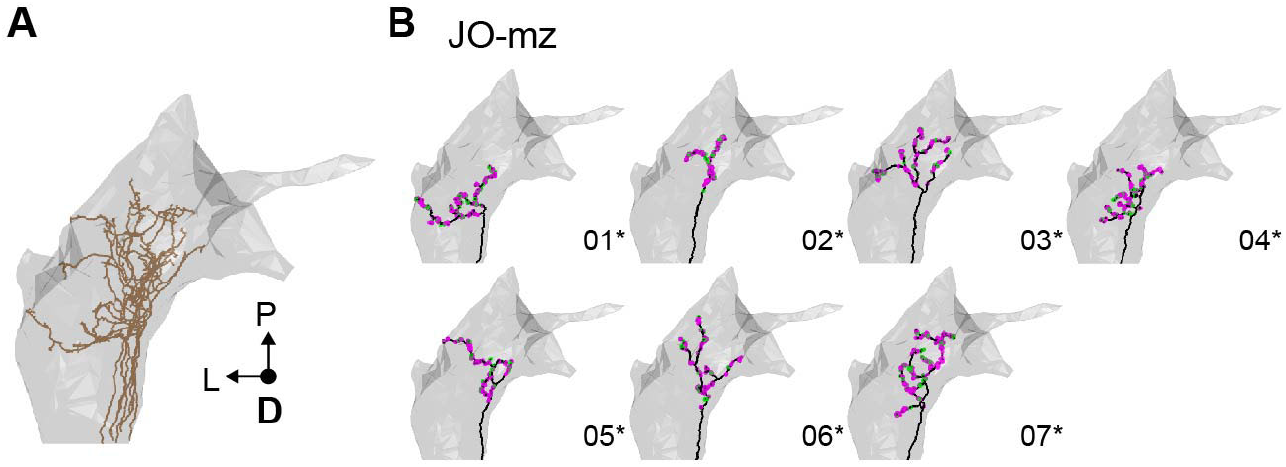
Individual reconstructed multiple zone-projecting JONs. (**A**) Dorsal view of the reconstructed JONs that project to multiple zones (described in **Figure 1G**). (**B**) Dorsal view of individual JO-mz neurons. Numbers below each image indicate the corresponding reconstructed neuron in the EM dataset (e.g. JO_mz_01). Asterisks by the numbers indicate the JONs that have been completely reconstructed, including pre- and postsynaptic sites. Pre- and postsynaptic sites are labeled in magenta and green, respectively.

**Figure 1 – figure supplement 7.**
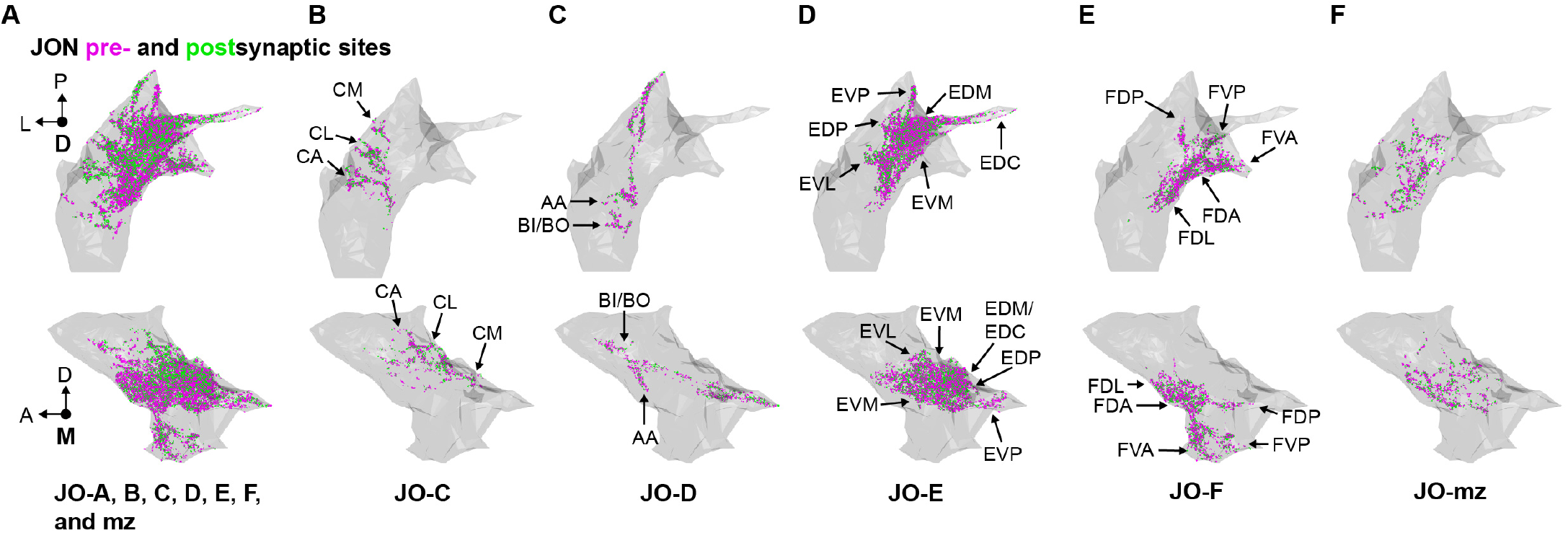
Distribution of JON synapses. (**A-F**) Shown are the medial (top) and dorsal (bottom) views of the JON synapses, subdivided into pre- and postsynaptic sites. All synapses of completely reconstructed JONs are shown in **A**. Synapses of zone C, D, E, F, or multiple zone (mz)-projecting JONs are shown in **B**, **C**, **D**, **E**, and **F**, respectively. JON pre- and postsynaptic sites are labeled in magenta and green, respectively. Zone subareas are indicated with labeled arrows.

**Figure 3 – figure supplement 1.**
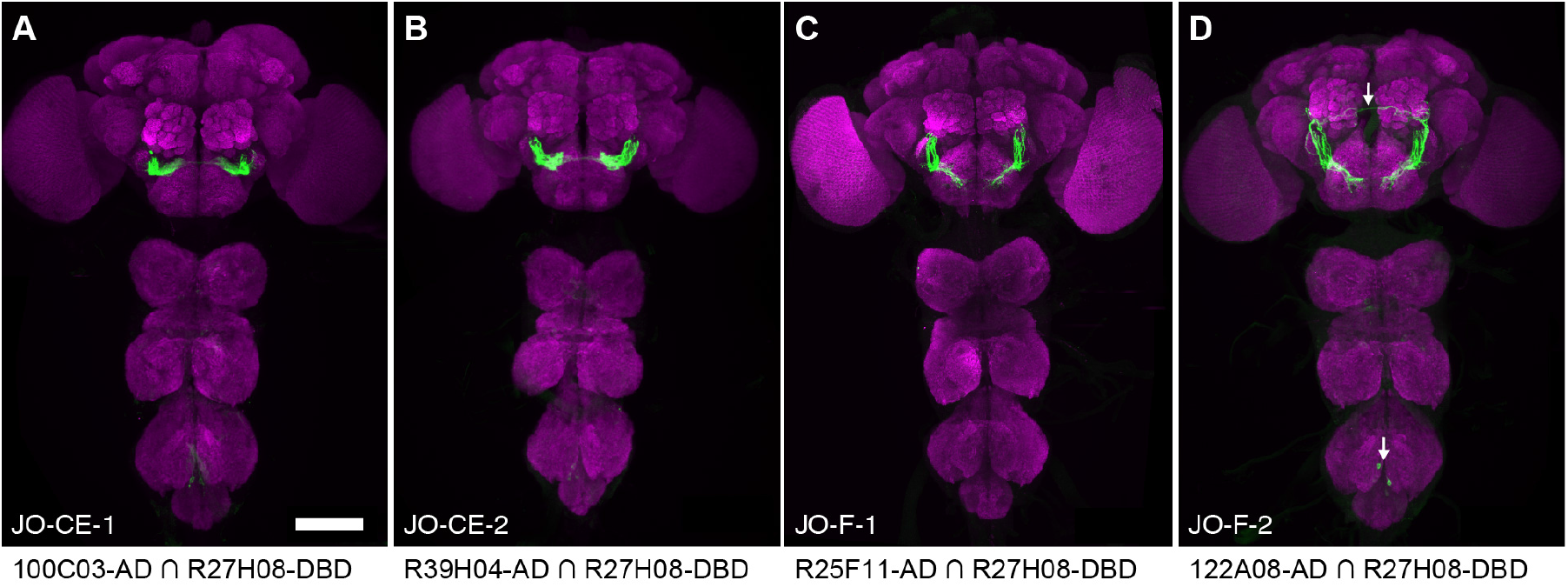
Driver lines that express in JO-C and -E or JO-F neurons. (**A-D**) Driver lines that express GFP in subpopulations of JONs. Shown are anterior views of the brains of JO-CE-1 (**A**), JO-CE-2 (**B**), JO-F-1 (**C**), and JO-F-2 (**D**). Images show maximum intensity projections of CNSs that are co-stained with anti-GFP (green) and anti-Bruchpilot (magenta). The arrows in **D** indicate neurons that are not JONs. Scale bar, 100 μm.

**Figure 3 – figure supplement 2.**
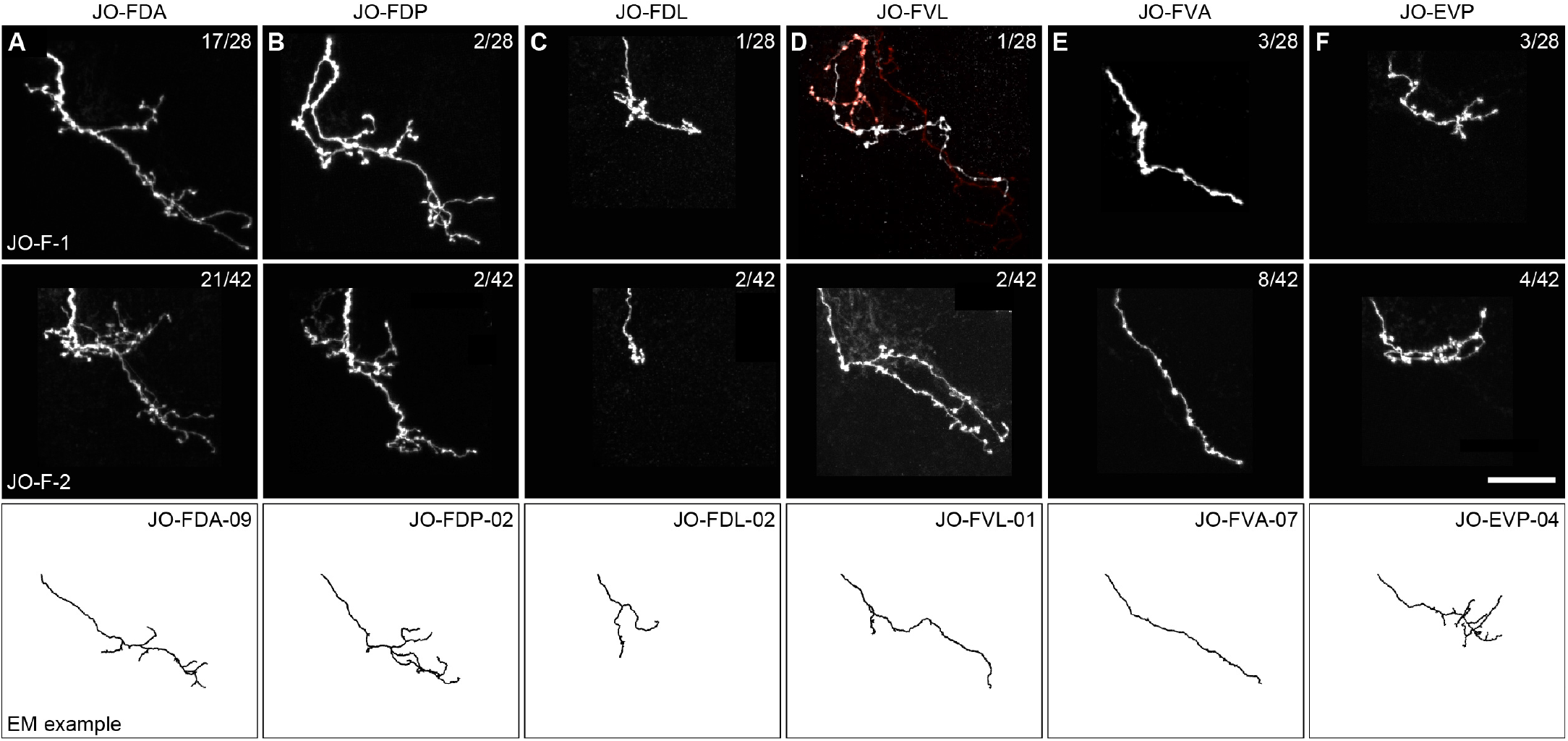
Stochastic labeling of individual JONs in the JO-F-1 and JO-F-2 expression patterns. (**A-F**) Anterior view of different MCFO-labeled JON types in the expression patterns of JO-F-1 (top) and JO-F-2 (middle). Shown are maximum intensity projections of each JON expressing a tagged protein that is stained using a tag-specific antibody (see Materials and methods for information about the tagged protein and antibodies used). Top right corner of each panel indicates the number of JONs that were labeled for each type versus the total MCFO-labeled JONs that we obtained (white neurons are the type indicated). Note that the number of individual labeled JONs does not add up to the total number of MCFO-labeled JONs (27 out of 28 for JO-F-1 and 39 out of 42 for JO-F-2). The additional neurons were not included in the analysis because they had ambiguous morphology that could not be definitively linked to a particular JON type. Bottom panels show EM reconstructed examples of each neuron type (neuron names are indicated in the top right corner). The JON types are JO-FDA (**A**), -FDP (**B**), -FDL (**C**), -FVL (**D**), -FVA (**E**), and -EVP (**F**). Scale bar, 20 μm.

**Figure 5 – figure supplement 1.**
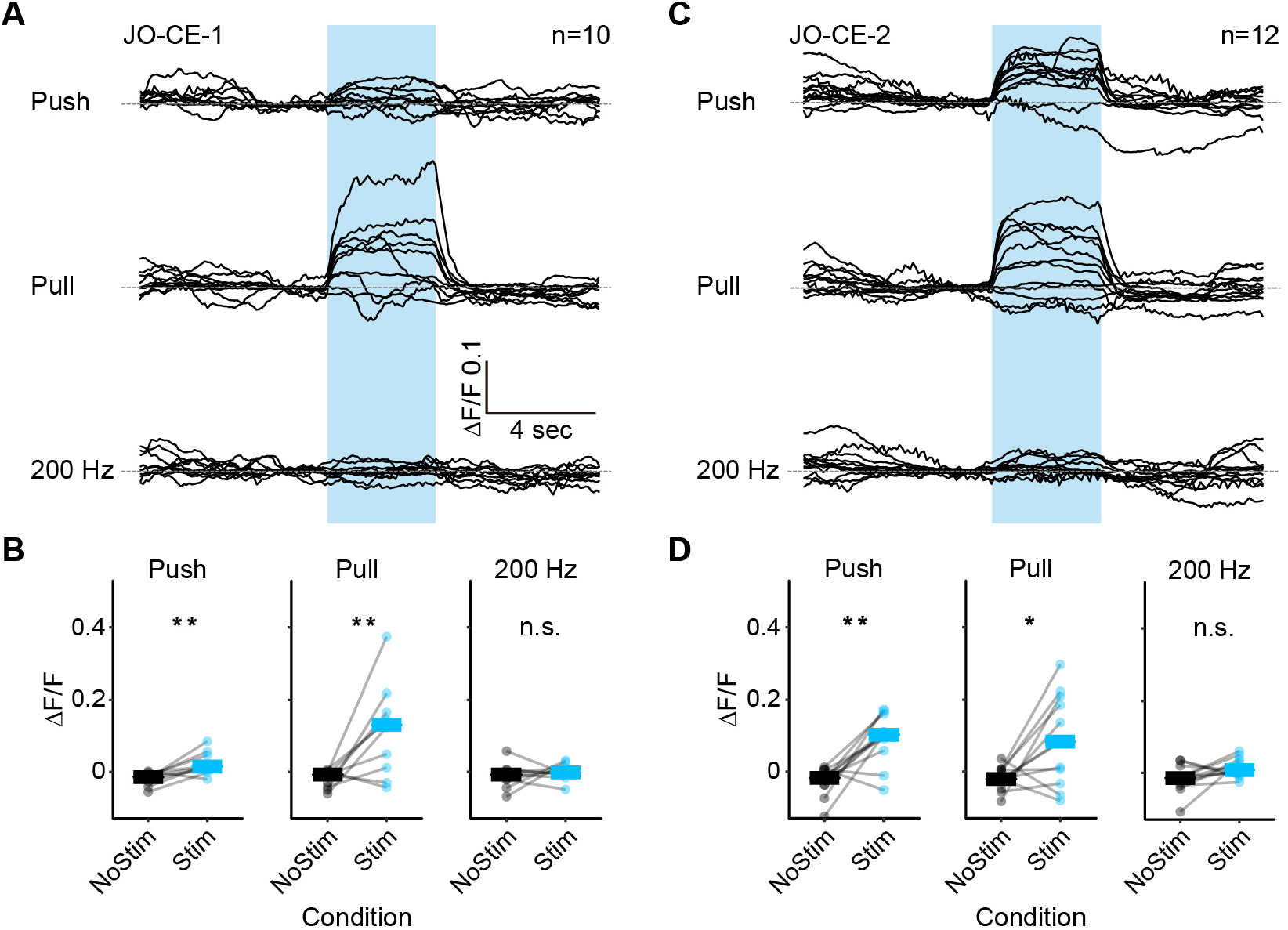
Responses of the JO-CE neurons to mechanical stimuli: raw traces and statistical analysis. (**A-D**) gCaMP6f was expressed in either JO-CE-1 (**A,B**) or JO-CE-2 (**C,D**). (**A,C**) Shown are the gCaMP6f fluorescence traces of individual trials while different mechanosensory stimulations were delivered (average traces are shown in **Figure 5B,C**). The stimulus duration was 4 seconds and is indicated with blue boxes. (**B**) Plots of the measured fluorescence before and during each stimulus (NoStim and Stim, respectively) for each trial (dots), and the average of the trials (bars). The Wilcoxon signed-rank test was used for each condition (asterisks indicate *p < 0.05, **p < 0.001).

**Figure 5 – figure supplement 2.**
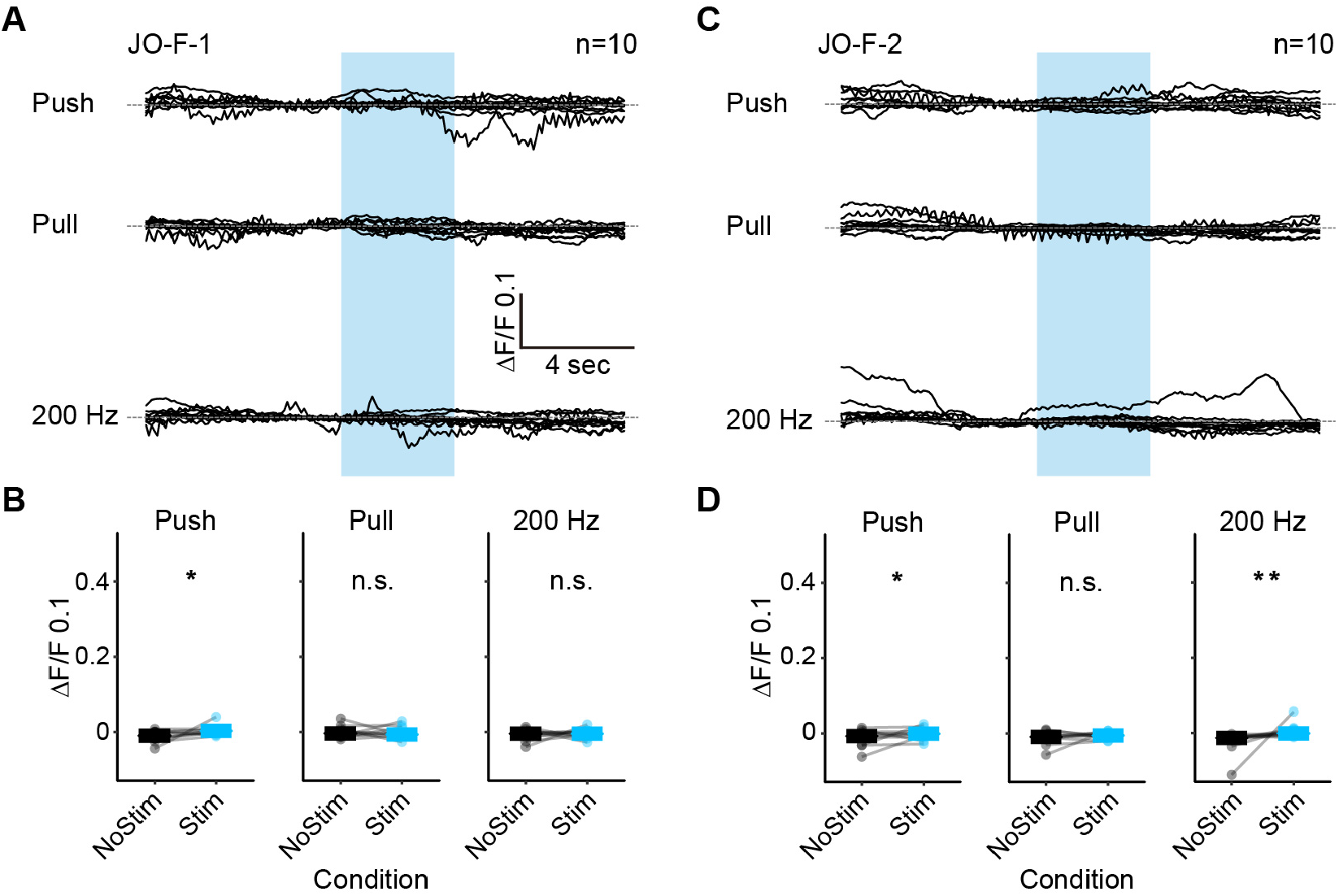
Responses of the JO-F neurons to mechanical stimuli: raw traces and statistical analysis. (**A-D**) gCaMP6f was expressed in either JO-F-1 (**A,B**) or JO-F-2 (**C,D**). (**A,C**) Shown are the gCaMP6f fluorescence traces of individual trials while different mechanosensory stimulations were delivered (average traces are shown in **Figure 5D,E**). The stimulus duration was 4 seconds and is indicated with blue boxes. (**B**) Plots of the measured fluorescence before and during each stimulus (NoStim and Stim, respectively) for each trial (dots), and the average of the trials (bars). The Wilcoxon signed-rank test was used for each condition (asterisks indicate *p < 0.05, **p < 0.001).

**Figure 6 – figure supplement 1.**
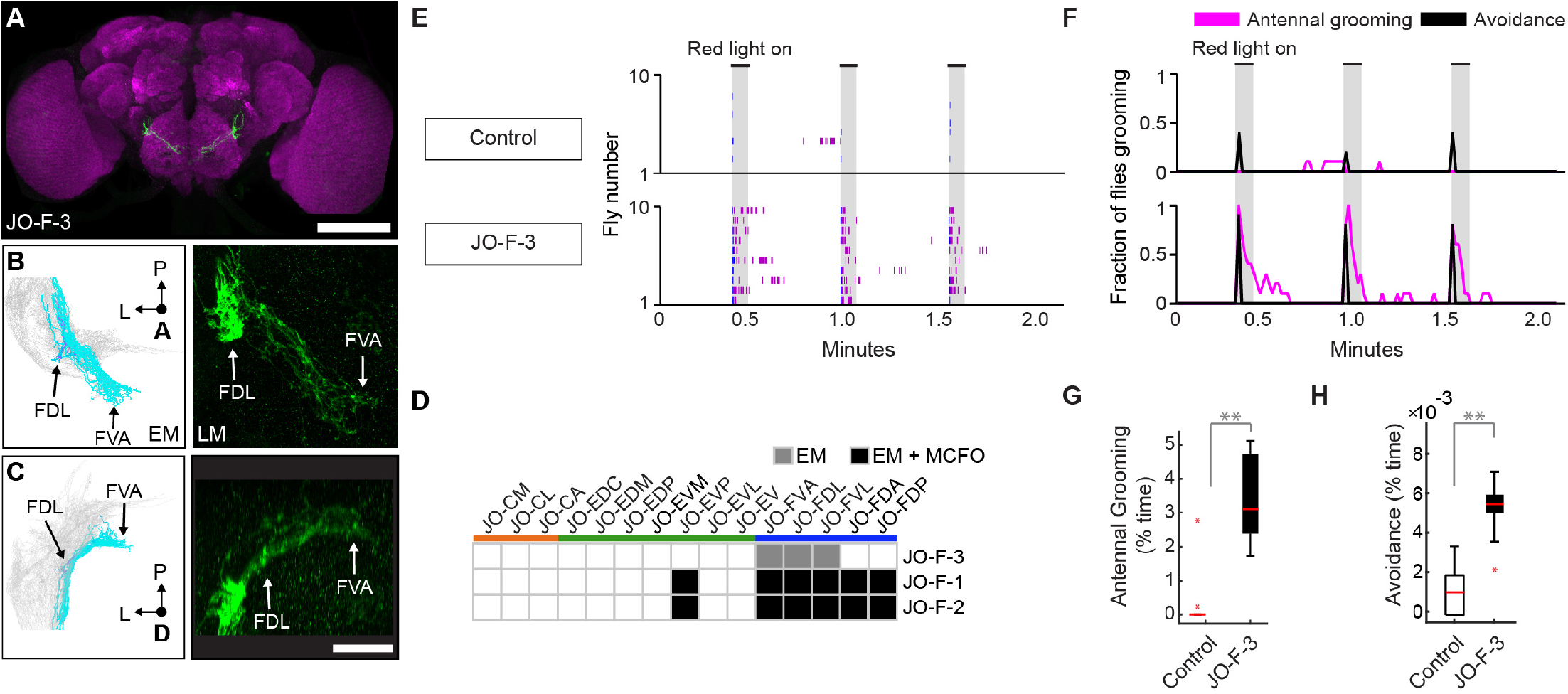
Driver line that expresses in JO-F neurons. (**A**) Shown is a maximum intensity projection of a brain (anterior view) of JO-F-3 (R60E02-LexA) expressing green fluorescent protein (myr::GFP). The brain is co-stained with anti-GFP (green) and anti-Bruchpilot (magenta). Scale bar, 100 μm. (**B,C**) Anterior (**B**) and dorsal (**C**) views of EM reconstructed JO-F neuron types (left panels) that are predicted to be in the expression pattern of JO-F-3 (right panels). Subareas are indicated with arrows. Scale bar, 20 μm. (**G**) Table of JON types that are proposed to be in the JO-F-3 expression pattern, compared with JO-F-1 and −2. The shade of each box indicates whether the predictions are supported by EM reconstructions alone (gray), or by EM and MCFO data (black). (**E,F**) Ethograms (**E**) and histograms (**F**) of manually scored video show the behaviors elicited with red light induced optogenetic activation. Ethograms and histograms are shown as described in **Figure 6**. (**G**,**H**) Percent time flies spent performing antennal grooming (**G**) or avoidance (**H**) with optogenetic activation of JONs targeted by JO-F-3. Box plots are shown as described in **Figure 6**.

**Figure 8 – figure supplement 1.**
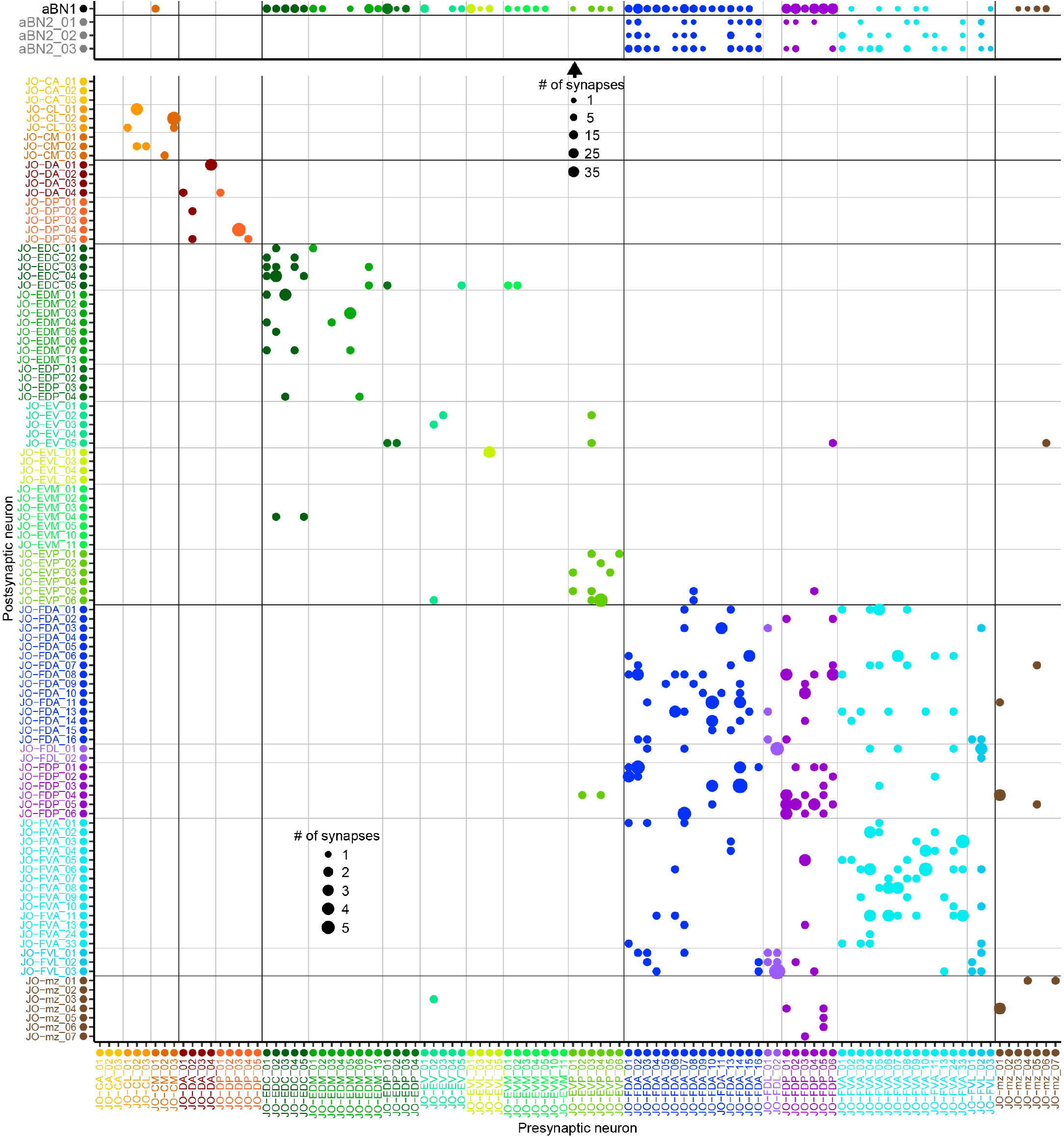
JON-to-aBN1/aBN2 and JON-to-JON synaptic connectivity. Plotted is a matrix of the major synaptic connections of the completely reconstructed neurons from this study (presynaptic neurons – x axis, postsynaptic – y axis). The different sized dots on the grid indicate the strength (# of synapses) for each connection. Synapse strength reference dots are shown for the top grid (table with arrow) and the bottom grid (bottom left quadrant). The top matrix shows the synaptic connections of the JO-C (orange), -E (green), -F (blue), and -mz (brown) neurons onto aBN1 (connection strength between 1 and 35), and the JO-F neuron connections to the aBN2 neurons (connection strength 1 to 7). Connections from JONs to aBN1/aBN2 neurons are axodendritic with the exception of 19 synapses onto the aBN1 axon (shown in **Figure 8 – supplement 3A**). The bottom matrix shows that JONs of the same type are almost exclusively presynaptic to each other, with connection strengths ranging from 1 to 5 synapses (all connections axo-axonic). The 4 aBN1-to-JON synapses and 4 synapses between aBN2 neurons and aBN1 are not shown. For a complete all-to-all connectivity table see **Supplementary file 2**.

**Figure 8 – figure supplement 2.**
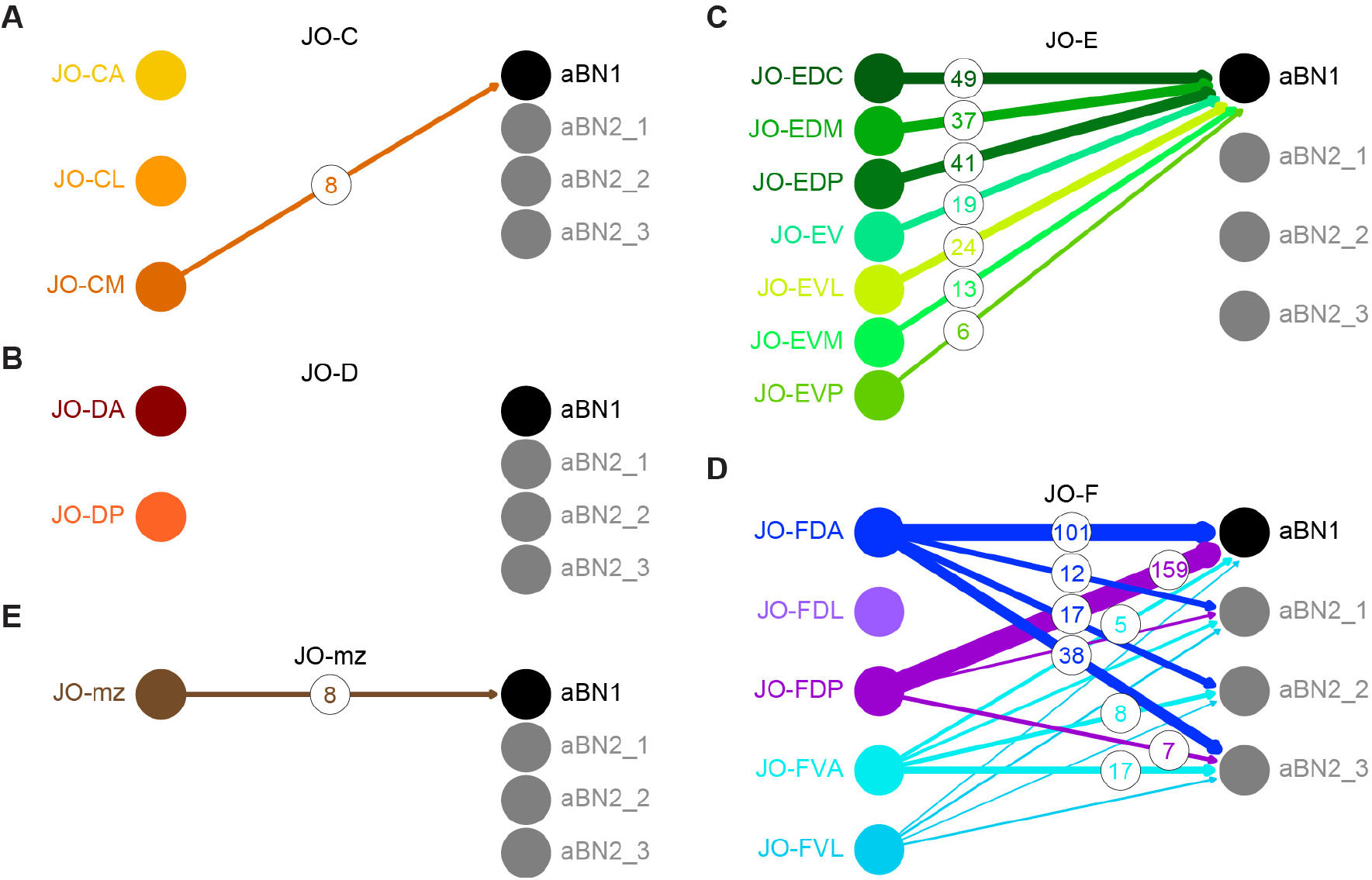
JON-to-aBN1 or aBN2 synaptic connections of fully reconstructed neurons from this study. Shown are the synaptic connections of the different JON types to aBN1 and aBN2 neurons for each subpopulation, including JO-C (**A**), of JO-D (**B**), JO-E (**C**), JO-F (**D**), and JO-mz (**E**). Connections from JONs to aBN1 or aBN2 neurons are axo-dendritic with the exception of 19 synapses onto the aBN1 axon (**Figure 8 – supplement 3A**). Arrow thickness represents number of synaptic connections. Numbers in circles indicate the number of synapses for connection strengths of more than 4.

**Figure 8 – figure supplement 3.**
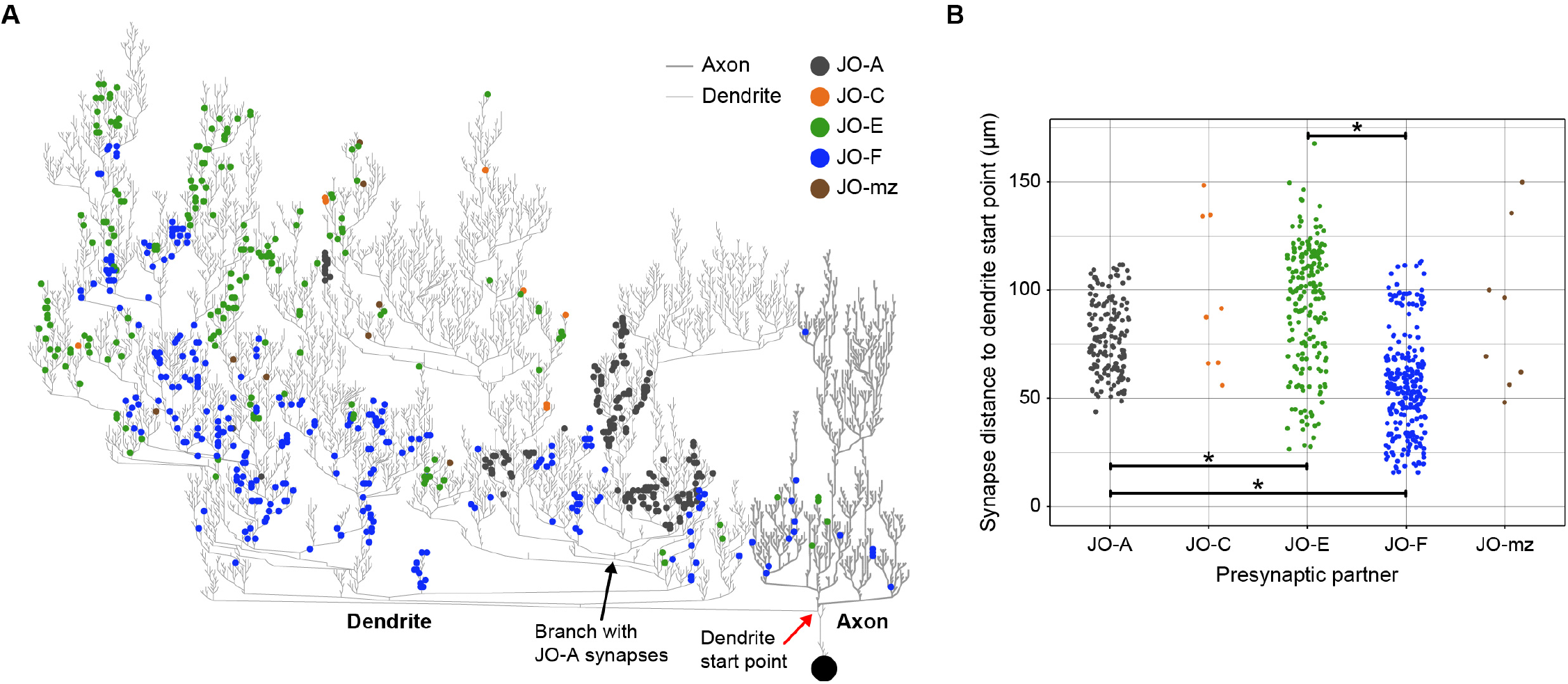
Distribution of JON synapses in the aBN1 dendrite: dendrogram. (**A**) Dendrogram of reconstructed aBN1 with the synapse locations of the different JON subpopulations. The JONs have 610 synapses onto the aBN1 dendrite (thin lines, left) and 19 synapses onto the aBN1 axon (thick lines, right). Synapses from JO-A (gray), -C (orange), -E (green), -F (blue) and -mz (brown) to aBN1 are indicated by colored dots. The JO-A neurons synapse onto a specific subset of aBN1 branches, whose convergence point is indicated with a black arrow. (**B**) Plot of distances of the different JON subpopulation postsynaptic sites to the start point of the aBN1 dendrite. This start point was defined as the point of bifurcation of the aBN1 primary neurite into axon and dendrite (indicated with a red arrow in **A**). JON-to-aBN1 synaptic sites are distributed along the proximal-to-distal axis of the aBN1 dendrite that can be measured as the distance from the dendritic start point. JO-E neuron synapses (green) are distributed in more distal dendritic space at distances that are significantly different from JO-A (gray, p = 1.35 x 10^-8^) and JO-F (blue, p = 2.6 x 10^-35^) neuron synapses. JO-F synapses are located more proximal to JO-A neuron synapses (p = 1.4 x 10^-24^) on the aBN1 dendrite. JON-to-aBN1 synapses located on the axon were excluded in **B**. Distributions of JO-C and -mz synaptic locations are shown but were excluded from the statistical analysis due to small connection strength (8 synaptic locations each).

**Figure 8 – figure supplement 4.**
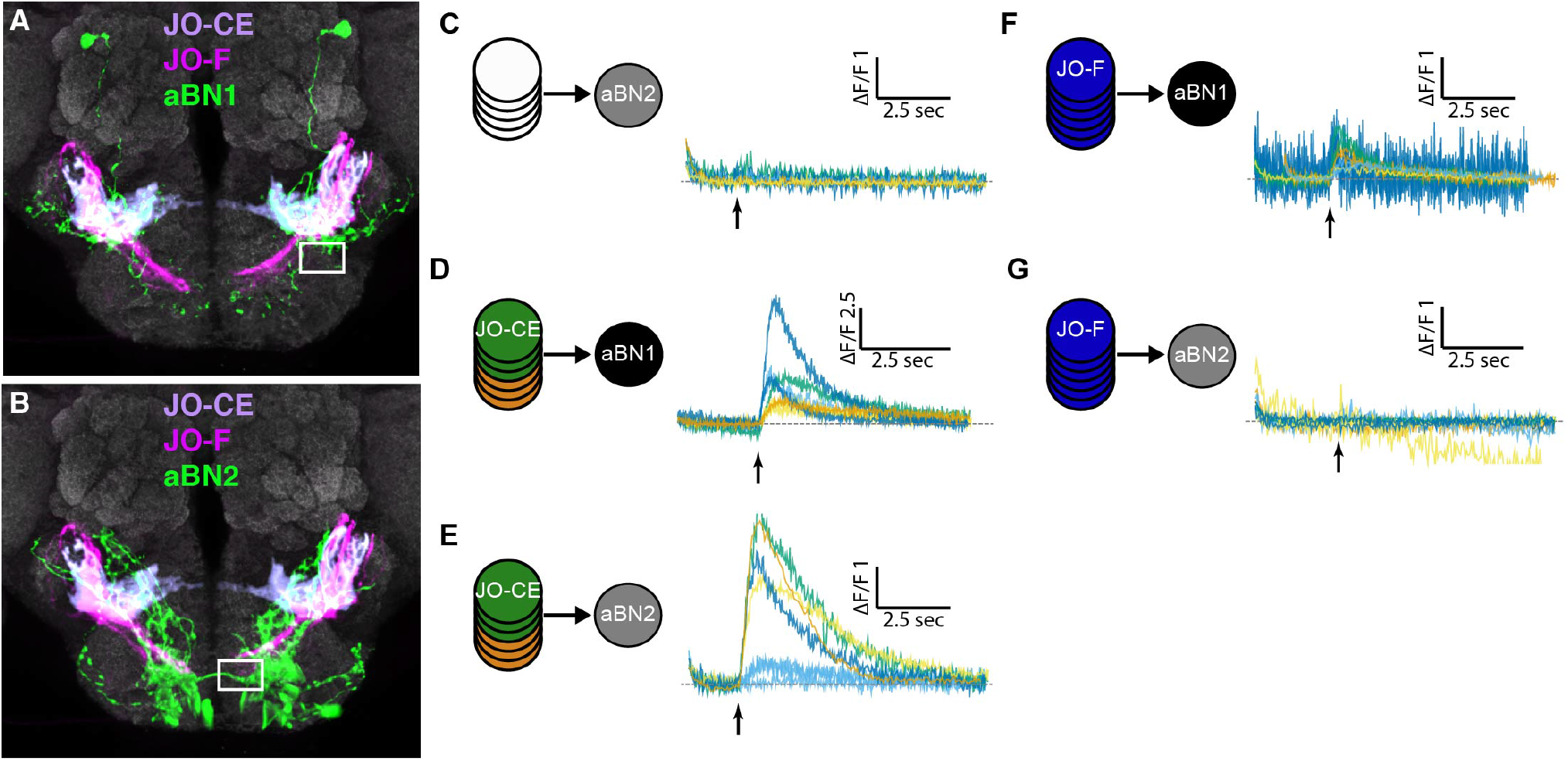
Functional connectivity experimental controls and raw data. (**A,B**) Representative examples of the regions of interest (ROIs, white boxes) that were selected for imaging fluorescence changes in aBN1 (**A**) and aBN2 (**B**). The JO-CE (purple) and -F neurons (magenta) are computationally aligned to show how they do not overlap with the ROIs. (**C**) Individual traces for the control experiment that shows no change in GCaMP6 fluorescence under our red light stimulus conditions in the absence of CsChrimson expression in JONs. Red light intensity was set at 500 μW/mm^2^. 30 light pulses were delivered, where each light pulse was 2 ms long, and the interpulse intervals were 18 ms (5 flies tested). Each trace shown is the average of 4 responses recorded at ~20 second intervals. (**D-G**) Individual traces of GCaMP6 responses in aBN1 or aBN2 neurons with CsChrimson-mediated JO-CE and -F neuron optogenetic activation (average traces shown in **Figure 8I**). Each tested neuronal pair is shown with labeled circles, including JO-CE to aBN1 (**D**), JO-CE to aBN2 (**E**), JO-F to aBN1 (**F**), and JO-F to aBN2 (**G**). Red light intensity was set at 50 μW/mm^2^ (**D,E**) and 500 μW/mm^2^ (**F,G**). 20 light pulses were delivered, where each light pulse was 2 ms and the interpulse intervals were 18 ms. Multiple runs were sometimes performed for a given CNS (shown by the same colored traces in a row). At least five flies were tested for each pair. See Materials and methods for driver lines used for each tested pair.

**Figure 8 – figure supplement 5.**
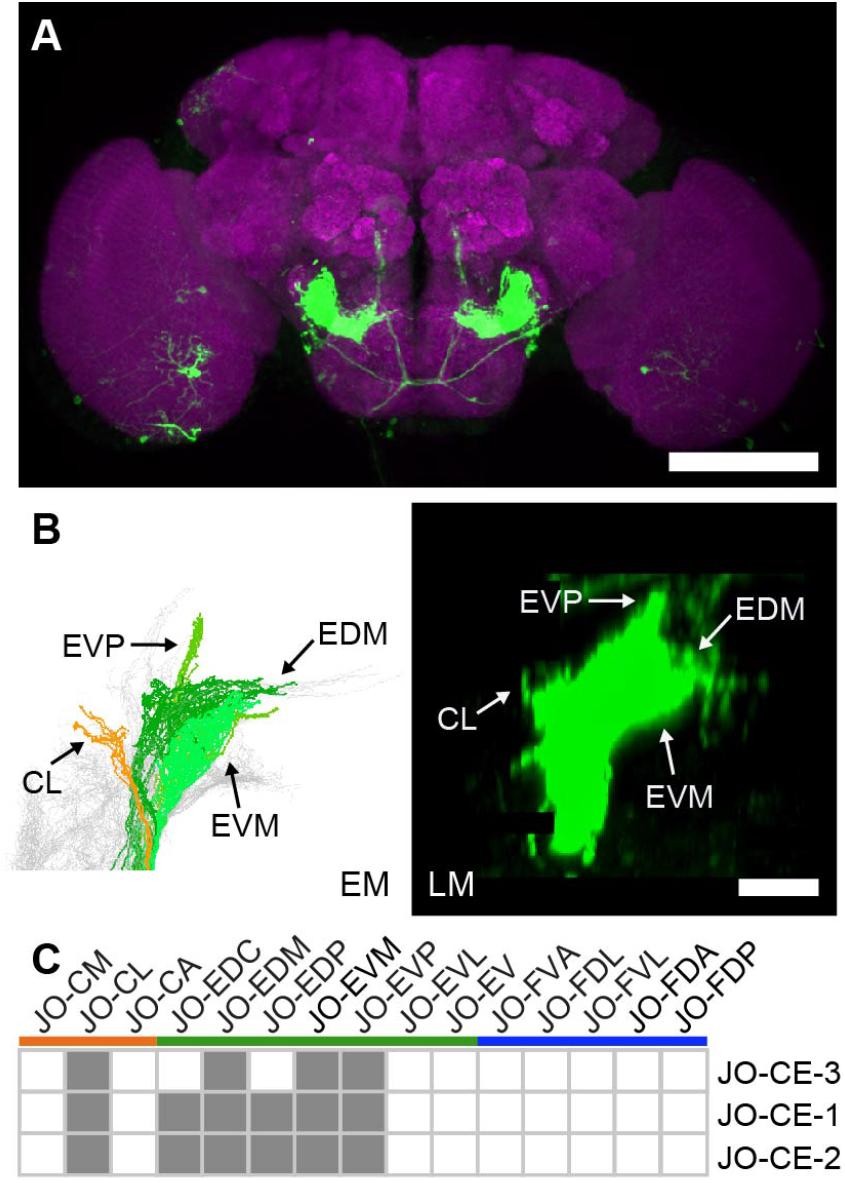
LexA driver line that expresses in JO-CE neurons. (**A**) Shown is a maximum intensity projection of a brain (anterior view) of JO-CE-3 (100C03-LexA) expressing green fluorescent protein (myr::GFP). The brain is co-stained with anti-GFP (green) and anti-Bruchpilot (magenta). Scale bar, 100 μm. (**B**) Dorsal view of EM reconstructed JO-CE neuron types (left) that are predicted to be in the expression pattern of JO-CE-3 (right). Subareas are indicated with arrows. Scale bar, 20 μm. (**C**) Table of JON types that are proposed to be in the JO-CE-3 expression pattern, compared with JO-CE-1 and −2. The shaded boxes indicate predictions that are supported by EM reconstructions. This line was used for csChrimson expression in the JO-CE neurons for testing functional connectivity with aBN1 and aBN2 neurons in **Figure 8I**.

**Video 1. EM reconstructed JONs.**

**Video 2. Optogenetic activation of JO-CE neurons elicits antennal grooming and wing flapping.**

**Video 3. Optogenetic activation of JO-F neurons elicits antennal grooming and avoidance.**

**Video 4. Optogenetic stimulus induces avoidance in control flies.**

**Video 5. EM reconstructed JON, aBN1, and aBN2 neurons and their synapses.**

